# Modeling CSF circulation and the glymphatic system during infusion using subject specific intracranial pressures and brain geometries

**DOI:** 10.1101/2024.04.08.588508

**Authors:** Lars Willas Dreyer, Anders Eklund, Marie Elisabeth Rognes, Jan Malm, Sara Qvarlander, Karen-Helene Støverud, Kent-Andre Mardal, Vegard Vinje

## Abstract

**Background:** Infusion testing is an established method for assessing CSF resistance in patients with idiopathic normal pressure hydrocephalus (iNPH). To what extent the increased resistance is related to the glymphatic system is an open question. Here we introduce a computational model that includes the glymphatic system and enables us to determine the importance of 1) brain geometry, 2) intracranial pressure and 3) physiological parameters on the outcome of and response to an infusion test.

**Methods:** We implemented a seven-compartment multiple network porous medium model with subject specific geometries from MR images. The model consists of the arterial, capillary and venous blood vessels, their corresponding perivascular spaces, and the extracellular space (ECS). Both subject specific brain geometries and subject specific infusion tests were used in the modeling of both healthy adults and iNPH patients. Furthermore, we performed a systematic study of the effect of variations in model parameters.

**Results:** Both the iNPH group and the control group reached a similar steady state solution when subject specific geometries under identical boundary conditions was used in simulation. The difference in terms of average fluid pressure and velocity between the iNPH and control groups, was found to be less than 6 % during all stages of infusion in all compartments. With subject specific boundary conditions, the largest computed difference was a 75 % greater fluid speed in the arterial perivascular space (PVS) in the iNPH group compared to the control group. Changes to material parameters changed fluid speeds by several orders of magnitude in some scenarios. A considerable amount of the CSF pass through the glymphatic pathway in our models during infusion, i.e., 28% and 38% in the healthy and iNPH patients, respectively.

**Conclusions:** Using computational models, we have found the relative importance of subject specific geometries to be less important than individual differences in terms of fluid pressure and flow rate during infusion. Model parameters such as permeabilities and inter-compartment transfer parameters are uncertain but important and have large impact on the simulation results. The computations predicts that a considerable amount of the infused volume pass through the brain either through the perivascular spaces or the extracellular space.

## Introduction

Idiopathic Normal Pressure Hydrocephalus (iNPH) is a partially reversible form of dementia characterised by enlarged ventricles. Typical symptoms are gait disturbance, urinary incontinence, and cognitive decline which may improve after shunt treatment Malm and Eklund [2006a]. Infusion tests have been shown to be an effective procedure for predicting if a patient is going to respond well to treatment Kahlon et al. [2005], Toma et al. [2013]. An infusion test measures the outflow resistance, *R*_*out*_, of the cerebrospinal fluid system as a whole. This resistance is in general significantly larger in iNPH patients than in healthy individuals Kahlon et al. [2005], Qvarlander et al. [2014]. Additional quantities like compliance, time required to reach pressure equilibrium, and craniospinal pressure volume index (PVI) are also common indicators Eklund et al. [2007], Wåhlin et al. [2010].

In 2012 Iliff et al. [2012] the glymphatic pathway for cerebrospinal fluid flow through the murine brain was identified and suggested to be important for clearance of metabolic waste. Accumulation of metabolic waste is common in dementia Turner [2021], Jack Jr et al. 2018] and is suggested to be caused by a malfunctioning glymphatic pathway. The glymphatic system and its pathways have been detailed in mice. Here, evidence suggest that bulk CSF flow occurs in pial PVS Mestre et al. [2018], Raghunandan et al. [2020], Abbott et al. [2018] and these flow velocities are around 20 *μ*m/s. Bulk flow of interstitial fluid has been estimated in the range from 10 nm/s Holter et al. [2017], Hladky and Barrand [2016, 2022] to 1.7 *μ*m/s Ray et al. [2019], Rosenberg et al. [1980]. In humans, less is know about the glymphatic pathways. Here, iNPH patients are particularly interesting as the relation between outflow resistance and pressure during infusion tests is well characterized in this patient group. Furthermore, the CSF dynamics of the brain is significantly altered in patients suffering from iNPH. Pulsatile flow of CSF in the aqueduct of Sylvus is larger in iNPH patients than in healthy individuals. Furthermore, the net flow is potentially retrograde rather than antegrade Lindstrøm et al. 2019], Eide et al. [2021]. Finally, tracer intrathecally injected into the brain has a significantly delayed clearance rate in iNPH patients Eide and Ringstad [2019], Ringstad et al. [2018], Eide et al. [2021]. If untreated, iNPH might lead to irreversible damage to brain tissue, and hence early detection and intervention is crucial Andrén et al. [2021]. Infusion testing is both a reliable and frequently used method for selecting patients for surgery Kahlon et al. [2005], Malm and Eklund [2006b], Peterson et al. [2016]. There exist several computational modeling studies Sobey and Wirth [2006], Wirth and Sobey [2006], Tully and Ventikos [2011], Dutta-Roy et al. [2008] of hydrocephalus and iNPH which predate the the glymphatic system by Illiff et al.Iliff et al. [2012].

These studies only model the interstitial space and do not include the perivascular pathways. To consider the glymphatic pathway, Vinje et al. Vinje et al. [2020] constructed a 0D multi-compartmental model investigating how fluid flow patterns change in the brain during infusion tests. Furthermore, Guo et al. Guo et al. [2020] utilized a model similar to ours to study both the glymphatic pathway and subject specific geometries to model cerebral CSF dynamics, but they did not consider infusion tests. Finally, we mention that the mathematical theory for such systems has recently been studied in several papers, see Lee et al. Lee et al. [2019] for an overview. However, so far, the relative importance between 1) subject specific geometries 2) intracranial pressure and 3) physiological parameters has not been assessed by computational models.

In this study we explored the differences between iNPH patients and healthy controls during an infusion test in terms of fluid pressure, speed and flow in the CSF and Interstitial Fluid (ISF). In 47 subject specific geometries, we studied fluid pressure in the perivascular spaces (PVS) and the extracellular space (ECS) of the brain, as well as ECS water transport to blood networks and blood perfusion. First, we tested whether the difference in geometry alone was sufficient to obtain differences in intracranial pressure (ICP) between the two groups during infusion. Second, we investigated if subject specific boundary conditions, in terms of subarachnoid CSF pressure and arterial inflow, are important for our model. Finally, we investigated if changes to brain physiology and the glymphatic pathway may play a role in the iNPH response to infusion. These changes included variations to permeability in the ECS and PVS, and variations in fluid flow pathways between the PVS and ECS.

## Methods

### Subject data and mesh generation

From previous studies Malm et al. [2011], Jacobsson et al. [2018] we obtained T1-weighted MRI images (turbo field echo (T1W-TFE) sequence) and pressure measures of 47 subjects (33 healthy and 14 iNPH patients), see table 1. MR images was obtained from a 3T Philips Achieva scanner (Philips Healthcare, Best, the Netherlands) with resolution 1.0 *×* 1.0 *×* 1.0 mm, which was interpolated to get an image with resolution 0.3 *×* 0.3 *×* 1.0 mm. Both the iNPH patients and the controls underwent an infusion test where mock CSF is injected into the lumbar canal. As the CSF volume increases, the ICP rises and parameters such as *R*_*out*_ are measured Andersson et al. [2005], Wåhlin et al. [2010].

**Table 1:** Characteristic data and range for the iNPH and control cohorts Qvarlander et al. [2017]. Table shows average value with one standard deviation. The reference pressure *p*_*ref*_ is the steady state pressure at infinite compliance Avezaat and van Eijndhoven [1986].

| Parameters | Healthy ($N = 35$) | iNPH ($N = 16$) |
| --- | --- | --- |
| Sex (F/M) | 19/16 | 7/9 |
| Age (years) | $71 \pm 5$ (64-81) | $73 \pm 5$ (64-82) |
| Mini-Mental State Examination (MMSE) | $29 \pm 1$ (28-30) | $27 \pm 3$ (21-30) |
| Ventricular volume (ml) | $43 \pm 20$ (16-99) | $181 \pm 60$ (95-310) |
| Evans Index | $0.29 \pm 0.03$ (0.23-0.36) | $0.39 \pm 0.04$ (0.33-0.49) |
| Aqueduct cross-sectional area (cm<sup>2</sup>) | $0.10 \pm 0.02$ (0.09-0.16) | $0.23 \pm 0.17$ (0.09-0.77) |
| Heart rate (bpm) | $64 \pm 9$ (47-85) | $72 \pm 11$ (52-87) |
| Brain volume (l) | $1.04 \pm 0.10$ | $1.03 \pm 0.08$ |
| Pial surface area (dm<sup>2</sup>) | $20.5 \pm 1.75$ | $20.4 \pm 1.71$ |
| Ventricular surface area (dm<sup>2</sup>) | $1.89 \pm 0.43$ | $3.30 \pm 0.47$ |
| Outflow resistance $R_{out}$ (mmHg/(ml/min)) | $10.03 \pm 4.71$ | $18.06 \pm 7.84$ |
| Arterial inflow $B_{in}$ (ml/min) | $712.5 \pm 172.4$ | $653.4 \pm 172.2$ |
| Reference pressure $p_{ref}$ (mmHg) | $8.92 \pm 2.48$ | $9.26 \pm 3.08$ |

We considered the two groups both on subject specific level and group level. At the group level we created average images of the control and iNPH group, respectively. The images were preprocessed using Statistical Parametric Mapping software (SPM12; Wellcome Department of Cognitive Neurology, University College London, London, United Kingdom). First, the T1-images were segmented into grey matter, white matter, and CSF. Then, we used DARTEL Ashburner [2007] for image registration into group-specific templates (averages) of the control and iNPH group, respectively. The normalised images were finally aligned into MNI space and smoothed using a 1.0 mm full width at half maximum Gaussian filter.

We performed a Freesurfer Dale et al. [1999] segmentation on each of the subject specific images and on the average images. Based on the Freesurfer segmentation, we generated computational meshes which we used in the numerical simulations. Only the segmentations for the brain stem, and the grey and white matter in the cerebrum and cerebellum was used in the mesh generation process, and the meshes were generated using SVMTK Mardal et al. [2022], Valnes and Schreiner [2021], an axial slice of each mesh is shown in figure 1. One mesh was created for each patient, consisting of three subregions given by the segmented white matter, grey matter and brain stem.

**Figure 1.**
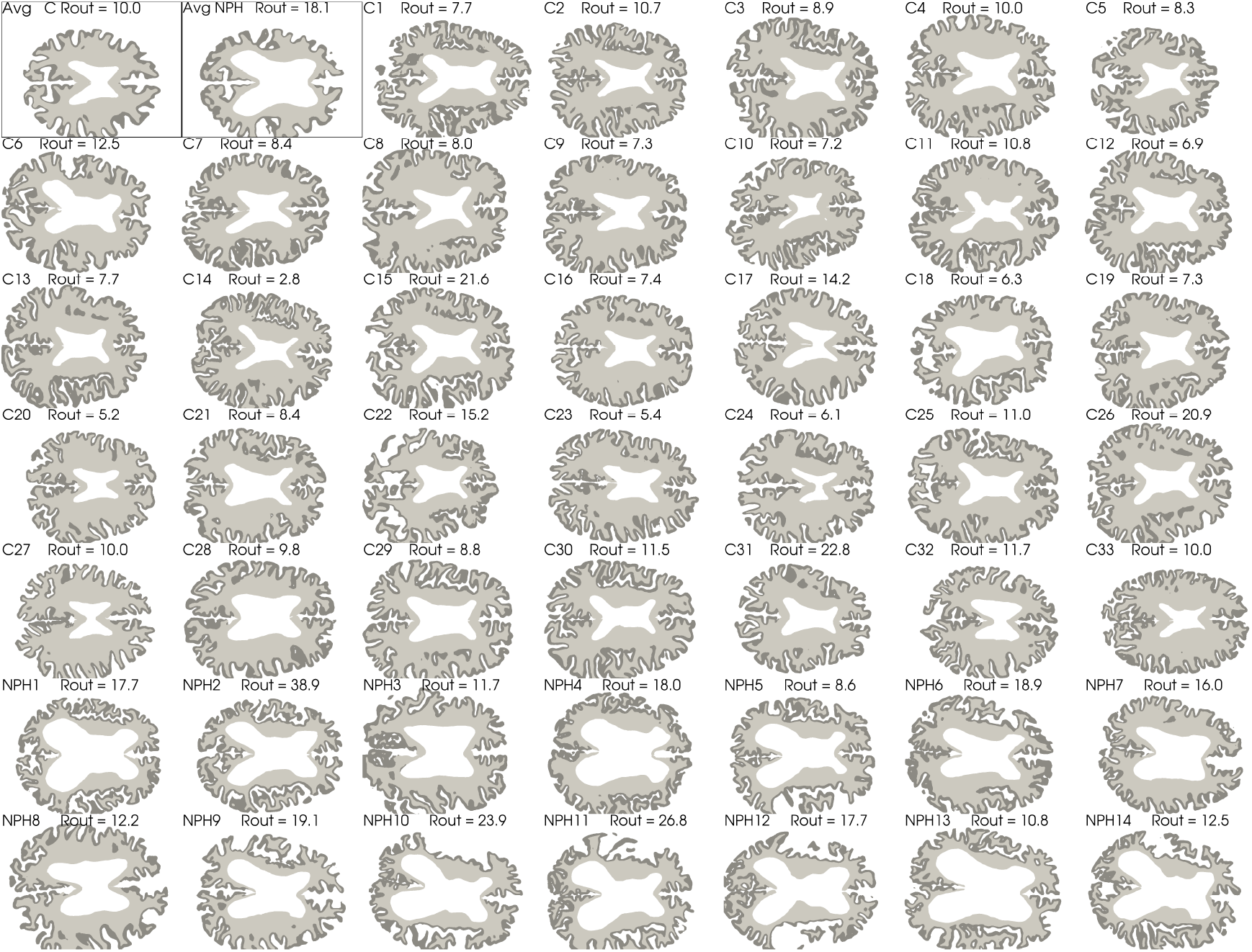
Axial slices of each subject as well as the average geometry marked as Avg C and Avg NPH for the control and iNPH groups respectively. The segmented grey matter is shown in dark grey, and the white matter is shown as a lighter shade of grey. The white space in the middle show the lateral ventricles. The two average geometries are shown first in the top left corner and are highlighted with a thin black frame.

### Governing Equations

We modelled the brain as a porous medium using a modified version of the MPET framework Tully and Ventikos [2011], where elastic deformations are ignored. Our model consists of seven compartments, namely arterial (a), capillary (c), and venous (v) blood compartments, their corresponding perivascular compartments (pa, pc, pv), and the extracellular space (e). Deformations of the brain parenchyma during an infusion were assumed to be negligible, yielding a simplified set of equations Dreyer [2022] given by

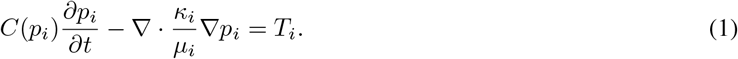

Here, *p*_*i*_ is the pressure in compartment *i*, while *C*(*p*_*i*_), *κ*_*i*_ and *μ*_*i*_ denotes the compartmental compliance, permeability and viscosity, respectively. The term *T*_*i*_ denotes the total fluid transfer between compartment *i* and its connected compartments and was modelled following Tully and Ventikos [2011], Guo et al. [2019] by:

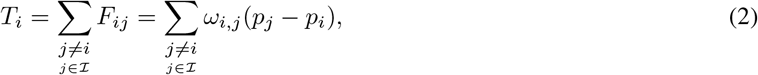

with *ω*_*i,j*_ and *F*_*ij*_ being the transfer coefficient and total fluid transfer between compartment *i* and *j* respectively. The pressure fields computed in the governing equations can be used to find the fluid pore velocity in each compartment,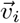, defined by

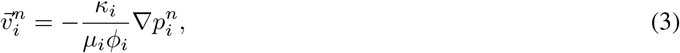

where *ϕ*_*i*_ is the compartmental porosity. The pore velocity is the velocity with which fluids travel through the individual pores, and is related to the Darcy velocity 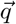 by 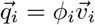. The Darcy velocity is defined as the volume flux through a pore divided by the pore’s cross sectional area. The model is illustrated graphically in figure 2.

**Figure 2.**
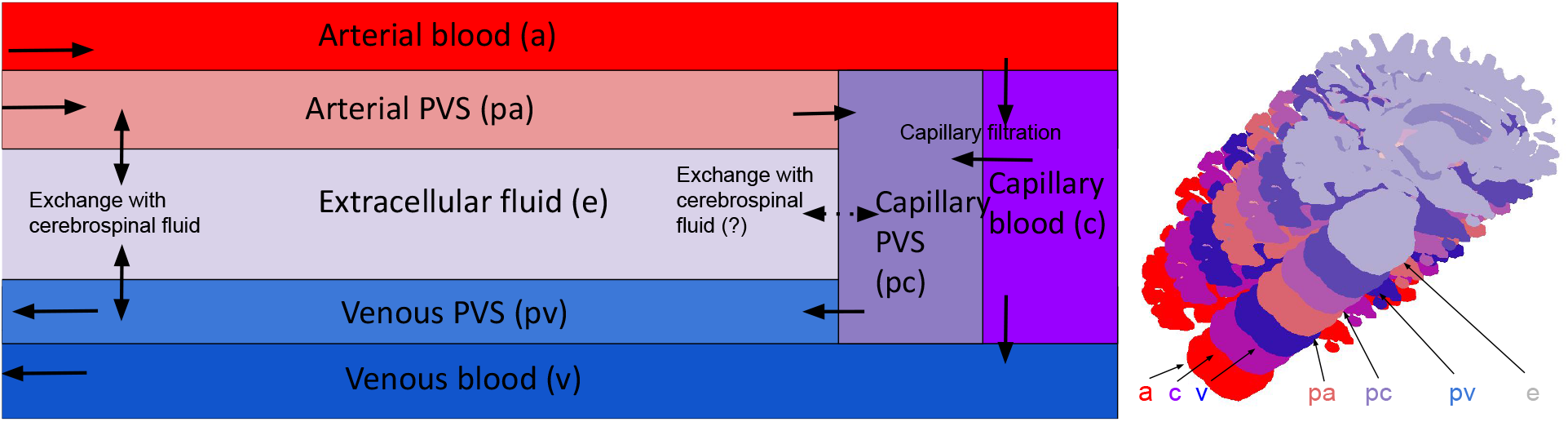
Conceptual sketch showing each compartment and their connections. Blood enters the brain through the arterial compartment and flows into the capillaries. From there, most of the blood flows further to the venous compartment and then out from the brain parenchyma, but a small fraction enters the perivascular space by means of capillary filtration. This is one of the two entryways for fluid into the PVS, the other being inflow through the arterial PVS. From there, the CSF can either keep flowing through the perivascular spaces, going first to the capillary PVS before travelling further to the venous PVS from which it leaves the parenchyma through pial sleeves alongside the cerebral veins. Alternatively, the CSF can flow through the ECS and to the venous PVS. In our base model, the possible fluid exchange between the ECS and capillary PVS was assumed to not be present, and is therefore marked with a dotted arrow. We remark that the focus here is on the compartments of the parenchyma and that the various exit routes are lumped together through one ordinary differential equations enforced uniformly at the brain surface.

### Parameters

In the governing equations, we use values from the literature to determine the permeabilities and porosities of each compartment as well as the intercompartmental transfer coefficient. Following Tully and Ventikos [2011], we set *μ*_blood_ = 3*μ*_CSF/ISF_, and the viscosity of CSF and ISF was set to the viscosity of water at 37 degrees centigrade. Finally, we assumed negligible deformations, and a low compliance *C*_*i*_ for all compartments. Larger vessels, namely arteries and veins were given larger compliance of *C*_a_ = *C*_v_ = 10^−4^ Pa^−1^, while the remaining five compartments have a compliance of *C*_*i*_ = 10^−8^ Pa^−1^.

Estimates for the flow resistance between different cerebral compartments are based on Vinje et al. [2020], whose model defined the intercompartmental transfer by

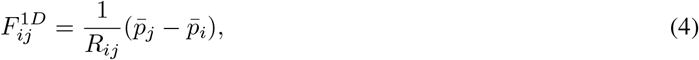

with 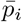 being the pressure in each 0D-compartment. Taking the volume average integral of equation 2, we can relate *ω*_*ij*_ to the one dimensional resistance *R*_*ij*_ by

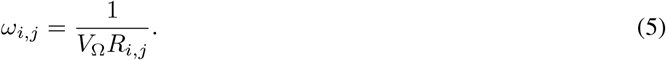

The transfer coefficients for flow between the PVS and the ECS, and the coefficients for flow between the PVS and capillary compartment were computed using the estimates of Vinje et al. [2020] for the one dimensional resistance *R*_*i,j*_.

We assumed the fluid transfer rate between blood networks to correspond to the flow and pressure drop observed experimentally. Hence, by equating equation 2 to the arterial inflow rate *B*_in_, we related the transfer coefficient between blood vessels to the expected pressure drop 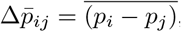,

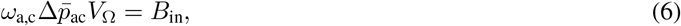

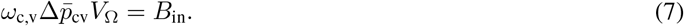

yielding We assumed a pressure drop of 60 mmHg from the arterial to the capillary compartment Paulson et al. [1990], Zagzoule and Marc-Vergnes [1986], and a pressure drop of 10 mmHg from the capillaries to the venous compartment Zagzoule and Marc-Vergnes [1986], Kinoshita et al. [2006]. The transfer coefficients depend on brain volume and arterial inflow, and hence we have listed the coefficients for a brain with *V*_Ω_ = 1 litre and *B*_*in*_ = 700 ml/min in table 2. These values were chosen to be similar to the data we used in our simulation, but both volume and blood flow varied between subjects and cohorts.

**Table 2:** Compartmental material parameters used in our base model. The porosities are dimensionless and are therefore marked with the units -.

| Parameter | Value | units | source |
| --- | --- | --- | --- |
| $\omega_{a,c}$ | $1.45 \cdot 10^{-6}$ | $\text{Pa}^{-1}\text{s}^{-1}$ | Zagzoule and Marc-Vergnes [1986] |
| $\omega_{c,v}$ | $8.75 \cdot 10^{-6}$ | $\text{Pa}^{-1}\text{s}^{-1}$ | Kinoshita et al. [2006] |
| $\omega_{c,pc}$ | $8.48 \cdot 10^{-10}$ | $\text{Pa}^{-1}\text{s}^{-1}$ | Vinje et al. [2020] |
| $\omega_{pa,e}$ | $1.86 \cdot 10^{-7}$ | $\text{Pa}^{-1}\text{s}^{-1}$ | Vinje et al. [2020] |
| $\omega_{pv,e}$ | $1.65 \cdot 10^{-7}$ | $\text{Pa}^{-1}\text{s}^{-1}$ | Vinje et al. [2020] |
| $\omega_{pa,pc}$ | $10^{-6}$ | $\text{Pa}^{-1}\text{s}^{-1}$ | Estimated |
| $\omega_{pc,pv}$ | $10^{-6}$ | $\text{Pa}^{-1}\text{s}^{-1}$ | Estimated |
| $\omega_{pc,e}$ | $10^{-10}$ | $\text{Pa}^{-1}\text{s}^{-1}$ | Estimated |
| $\kappa_a$ | $3.63 \times 10^{-14}$ | $\text{m}^2$ | Faghih and Sharp [2018], Vinje et al. [2020] |
| $\kappa_c$ | $1.44 \times 10^{-15}$ | $\text{m}^2$ | El-Bouri and Payne [2015] |
| $\kappa_v$ | $1.13 \times 10^{-12}$ | $\text{m}^2$ | Faghih and Sharp [2018], Vinje et al. [2020] |
| $\kappa_e$ | $2 \times 10^{-17}$ | $\text{m}^2$ | Holter et al. [2017] |
| $\kappa_{pa}$ | $3 \times 10^{-17}$ | $\text{m}^2$ | Vinje et al. [2020], Faghih and Sharp [2018] |
| $\kappa_{pc}$ | $1.44 \times 10^{-15}$ | $\text{m}^2$ | El-Bouri and Payne [2015] |
| $\kappa_{pv}$ | $1.95 \times 10^{-14}$ | $\text{m}^2$ | Vinje et al. [2020], Faghih and Sharp [2018] |
| $\phi_a$ | $1.09 \cdot 10^{-2}$ | - | Ito et al. [2001], Lee et al. [2001]. |
| $\phi_c$ | $2.31 \cdot 10^{-3}$ | - | Ito et al. [2001]. |
| $\phi_v$ | $1.98 \cdot 10^{-2}$ | - | Ito et al. [2001]. |
| $\phi_{pa}$ | $1.52 \cdot 10^{-2}$ | - | Mestre et al. [2018] |
| $\phi_{pc}$ | $2.31 \cdot 10^{-3}$ | - | Pizzo et al. [2018]. |
| $\phi_{pv}$ | $2.77 \cdot 10^{-2}$ | - | Mestre et al. [2018]. |
| $\phi_e$ | $1.40 \cdot 10^{-1}$ | - | Xie et al. [2013]. |

The resistance to fluid flow in each of our compartments from Vinje et al. [2020] can be converted to permeabilites for a three-dimensional compartment Dreyer [2022], Vinje et al. [2020]. If we let ∆:*p* denote the pressure change over a porous channel of length *L* and cross sectional area *A*, then the volume flux of a fluid with Darcy velocity *q* through this channel is given by

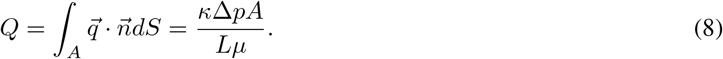

Following Vinje et al. [2020], we let *R*_*i*_ = ∆*p*_*i*_*/Q*_*i*_ denote the flow resistance in compartment *i*. We assumed homogeneity of the brain tissue, and that *L* and *A* are equal in all compartments. Using equation 8 with the definition of *R*_*i*_, the permeability and viscosity are related by

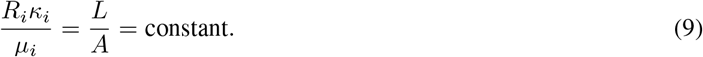

The permeability of the ECS, *κ*_*e*_, was estimated to be in the range spanning from 10 nm^2^ to 20 nm^2^ by Holter et al. [2017]. The authors do note that other estimates of the permeability reaches values in the order of 1000 nm^2^ to 4000 nm^2^ Morrison et al. [1994], Bobo et al. [1994], Prabhu et al. [1998], Basser [1992], but none of these latter references distinguish between perivascular and extracellular spaces. Hence, letting the extracellular permeability *κ*_e_ be 20 nm^2^, *μ*_ISF_ = *μ*_CSF_ = 0.75 mPa*·*s alltogether yields *L/A* = 1.21 *·* 10^−4^ m^−1^ according to Eq. (9).

In their computation of pericapillary resistance, Vinje et al. Vinje et al. [2020] considered two scenarios. One high resistance scenario where the capillary PVS was assumed to be 100 nm wide, based on the measurements of Bedussi et al. Bedussi et al. [2018]. These measurements were, however, conducted using fixation, which has been shown to shrink the PVS Mestre et al. [2018]. Furthermore, Pizzo et al. Pizzo et al. [2018] were able to image a part of the cerebral microvasculature of mice. Their image apparently show a PVS width of more than 1 *μ*m. Hence, we have in our model assumed the extended capillary gaps scenario from Vinje et al. Vinje et al. [2020] to be more representative of the actual physiology. We have listed the final values for each compartmental permeability in table 2.

The cerebral blood volume fraction was set at 3.3 % of the total brain volume, Perles-Barbacaru and Lahrech [2007], Ito et al. [2001], Lee et al. [2001] of which a third of the total cerebral blood volume (CBV) is arterial blood Ito et al. [2001], Lee et al. [2001]. We assumed a capillary blood fraction of 10 % of total CBV, with the remaining 57 % being venous blood Ito et al. [2001], Lee et al. [2001]. These volume fractions were used directly to determine the porosities for the vascular compartments. Furthermore, the perivascular porosities were determined as proportional to the porosities of the vascular compartments.

The perivascular porosities were computed as the relative size fraction between the PVS and its corresponding vasculature. We have for the arterial and venous perivascular spaces assumed the porosity to be 1.4 times that of the vasculature, as observed to be the area ratio on the pial surface in mice Mestre et al. [2018]. While the penetrating PVS might be smaller than the surface PVS, this proportionality gives a tangible upper bound on the porosities and hence a lower bound on the computed fluid velocities. For the capillary PVS, we used an ex-vivo image of the perivascular space of a cerebral microvessel Pizzo et al. [2018]. This image suggest a porosity proportionality of 1 between the pericapillary space and the capillaries. Finally, we have used an ECS porosity *ϕ*_*e*_ = 0.14, based on observations made of the murine brain Xie et al. [2013]. All porosities are listed in table 2.

The subarachnoid cerebrospinal fluid pressure, *p*_CSF_ was computed using the model of Vinje et al. [2020]:

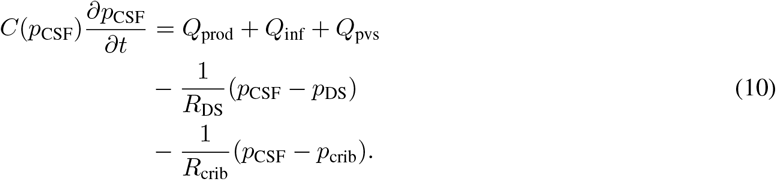

Here *p*_DS_ = 8.4 mmHg and *R*_DS_ = 10.81 mmHg/mL/min is the pressure and resistance at the dural sinus and *p*_crib_ = 0 (atmospheric pressure) and *R*_crib_ = 67 mmHg/mL/min is the cribriform plate pressure and resistance Vinje et al. [2020]. Furthermore, *Q*_inf_, *Q*_prod_ and *Q*_pvs_ denote the CSF in- and outflow from infusion, production in choroid plexus and outflow to the cerebral PVS respectively. The function *C*(*p*_*CSF*_) is the subarachnoid compliance, and was modelled following Vinje et al. [2020]:

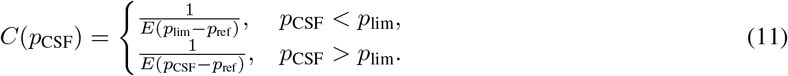

Here, *E* = 0.2 ml^−1^ is the elastance of the system, *p*_0_ = 13 mmHg is a lower threshold pressure and *p*_ref_ = 9 mmHg Vinje et al. [2020] in the base model. In cases where *p*_0_ or *p* were smaller than *p*_ref_, the reference pressure was set to be 1 mmHg less than *p*_0_ or *p*. We remark that *Q*_pvs_ was not included in Vinje et al. [2020], but was included here to ensure mass conservation.

The infusion test measurements from Qvarlander et al. [2017] were used to tune the boundary conditions of our computational model by fitting the best parameters of a lumped ordinary differential equation. In detail, we assumed the outflow resistances of the dural sinus and the cribriform plate, *R*_DS_ and *R*_crib_, to depend linearly on a real number *α*, i.e., 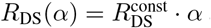 and 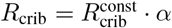. Here, the const-superscript indicates that it is a constant with a value equal to those used by Vinje et al. [2020]. We computed the pressure curves predicted by equation 10 for an infusion test with *Q*_inf_ = 1.5 ml/min for different values of *α*. Following Vinje et al. [2020], only *Q*_prod_ and *Q*_inf_ were included in these computations. Then, we computed the total *R*_out_ attained for each value of *α*. Using linear regression, we were able to then relate *R*_out_ to *α* and found the following relation to be the best fit

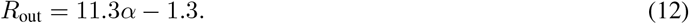

The mean squared error of the fit was 1.8 *·* 10^−4^ mmHg/(ml/min). Finally, the compliance *C*(*p*_CSF_) was tuned to depend on subject specific reference pressures *p*_ref_, modelling interpersonal variance in infusion response time.

### Simulation setup

Our domain was partitioned in three subdomains, namely grey matter, white matter and the brain stem. On the latter, homogeneous Neumann boundary conditions were used, ensuring no flow through the boundary of this part of the domain. The extracellular and pericapillary compartments were assumed disconnected from the subarachnoid CSF due to barriers such as pia and glia limitants, and had a homogenous Neumann condition on the entirety of the boundary surface. No compartment had fluid flow through more than one surface, and hence all had a no-flow condition on either the pial or ventricular surface. The boundary conditions which are not homogenous Neumann are listed in table 3. We enforced two sets of boundary conditions, generic and subject specific. These two cases differ in which values for arterial inflow *B*_in_, outflow resistance *R*_out_ and reference pressure *p*_ref_ were used. In the generic case, the average value in the control group was used for each of the aforementioned values. In the subject specific case, these parameters were set based on individual measurements.

**Table 3:** Table of each in- and outflow boundary conditions enforced on each compartment in the model. Here, *A*_pial_ is the pial surface area and *p*_CSF_ is the pressure in the subarachnoid space, and is defined in equation 10.

| Compartment | Boundary condition | Surface | Notes |
| --- | --- | --- | --- |
| Arterial | $\kappa_a \nabla p_a \cdot \hat{\mathbf{n}} = Q_{\text{in}}$ | Pial | $Q_{\text{in}} = B_{\text{in}}/A_{\text{pial}}$. |
| Capillary | $\kappa_c \nabla p_c \cdot \hat{\mathbf{n}} = -Q_{\text{prod}}$ | Ventricular | $Q_{\text{prod}} = 0.33 \text{ ml/min}$. |
| Venous | $\kappa_v \nabla p_v \cdot \hat{\mathbf{n}} = \beta_1 \left( \frac{p_{\text{DS}} + p_{\text{CSF}}}{2} - p_v \right)$ | Pial | $p_{\text{DS}} = 8.4 \text{ mmHg}$, $\beta_1 = 10^{-3}$. |
| Arterial PVS | $\kappa_{\text{pa}} \nabla p_{\text{pa}} \cdot \hat{\mathbf{n}} = \beta_2 (p_{\text{CSF}} - p_{\text{pa}})$ | Pial | $\beta_2 = 10^{-3}$. |
| Venous PVS | $\kappa_{\text{pv}} \nabla p_{\text{pv}} \cdot \hat{\mathbf{n}} = \beta_3 \left( \frac{p_{\text{CSF}} + p_{\text{DS}}}{2} - p_{\text{pv}} \right)$ | Pial | $\beta_3 = 10^{-7}$. |

The arterial and venous boundary conditions enforce a constant flow rate through the brain parenchyma, entering over the entire pial surface and leaving through veins penetrating the cortical surface. The boundary condition on the capillaries removed a fixed amount of fluid through the ventricular surface, modelling the fluid uptake by choroid plexi for CSF production. The perivascular boundary condition ensured that CSF flow through the PVS is driven by a pressure gradient between the PVS and SAS. On the venous PVS, the average of the Dural Sinus pressure and the subarachnoid CSF pressure is used to model the general increase in CSF pressure across the entire CNS, while the CSF still leaves the parenchyma alongside the cerebral veins. The parameters *β*_1_, *β*_2_ and *β*_3_ are numerical coefficients set to determine how easily the fluid can leave through the surface. Each simulation will run for a total length of *T* = 70 minutes. The simulation is run in three stages, with the first 16 minutes and 40 seconds being run without infusion to let the system reach steady state before turning on infusion. After this initial period, at *t* = 0 min, infusion starts at a constant rate of *Q*_inf_ = 1.5 ml/min. The infusion test lasts for 33 minutes and 20 seconds. Afterwards, when infusion stops the simulation runs for a period of 20 minutes to check for convergence back to the original steady state. The stage time length was set based on computed convergence time in terms of fluid velocity in a single subject.

The meshing software SVMTK uses a resolution parameter (RP) to determine an upper bound on cell size. Using numerical convergence testing, an RP of 32 (typically around 1.1M cells) was found to be sufficient for convergence with second order Lagrange polynomials as finite element basis functions, with a less than 2 % change in plateau pressure from a RP16 resolution (data not shown). The equations were discretised in time using a Backward-Euler scheme, and a time step of ∆*t* = 20*s* was sufficient to achieve convergence. (Determined graphically). All details regarding discretisation, mesh and time resolution can be found in Dreyer [2022], chapter 6. The simulations were performed using the Legacy FEniCS solver for python. Logg et al. [2012], Alnæs et al. [2015]

We performed eight sets of simulation experiments, listed in table 4. In the first three simulation sets, labelled as base model 1, 2 and 3, we explored the effect of geometric differences and differences in boundary conditions. Here, we used either subject specific or average geometries and either subject specific or generic boundary conditions. The difference in subject specific and generic boundary conditions lies in whether subject specific or the control group average value for *B*_in_, *p*_ref_ and *R*_*out*_ was used in equations 12, 10, 11 and 3. The generic boundary conditions used the average values for the control group for these parameters. The last set of simulations, labelled variation 1 to 5 investigated how changes to material parameters affected the response to infusion.

**Table 4:** Overview over the eight different sets of simulations that were performed in this study.

| Experiment | Geometry | BC | Other changes | # Simulations |
| --- | --- | --- | --- | --- |
| Base model 1 | Subject specific | Generic | None | 47 |
| Base model 2 | Subject specific | Subject specific | None | 47 |
| Base model 3 | Average | Subject specific | None | 47 |
| Variation 1 | Average | Generic | Pericapillary resistance set to either (in mmHg/ml/min):<br>$R_{pc} = 9.2 \cdot 10^{-3}$ (Base),<br>$R_{pc} = 3.32 \cdot 10^{-4}$ (Pizzo)<br>or $R_{pc} = 32.24$ (High) | 3 |
| Variation 2 | Average | Generic | CSF outflow resistances changed to (in mmHg/ml/min):<br>$R_{DS} = 10.81, R_{crib} = 67$ (Base),<br>$R_{DS} = 21.62, R_{crib} = 67$ (High),<br>$R_{DS} = 5.41, R_{crib} = 67$ (Low). | 3 |
| Variation 3 | Average | Generic | Changes to capillary filtration:<br>$\omega_{c,pc} = 8.48 \cdot 10^{-10} \text{ Pa}^{-1}\text{s}^{-1}$, no constant filtration (Base), or<br>$\omega_{c,pc} = 0 \text{ Pa}^{-1}\text{s}^{-1}$, with<br>constant filtration rate of 0.16 ml/min at ventricles (Alternate). | 2 |
| Variation 4 | Average | Generic | Perivascular transfer coefficients changed to (units $\text{Pa}^{-1}\text{s}^{-1}$):<br>$\omega_{pa,e} = 1.86 \cdot 10^{-7}, \omega_{e,pv} = 1.65 \cdot 10^{-7}, \omega_{pc,e} = 10^{-10}$ (Base),<br>$\omega_{pc,e} = 5 \cdot 10^{-7}$, the rest unchanged (Case 1),<br>$\omega_{pa,e} = 1.86 \cdot 10^{-6}, \omega_{e,pv} = 1.65 \cdot 10^{-6}, \omega_{pc,e} = 10^{-8}$ (Case 2),<br>$\omega_{pa,e} = 10^{-10}, \omega_{e,pv}$ unchanged, $\omega_{pc,e} = 1.86 \cdot 10^{-7}$ (Case 3),<br>$\omega_{pa,e}$ halved, $\omega_{e,pv}$ halved, $\omega_{pc,e}$ unchanged (Case 4). | 5 |
| Variation 5 | Average | Generic | Changes to extracellular permeability (Units $\text{m}^2$):<br>$\kappa_e = 2.0 \cdot 10^{-17}$ (Base),<br>$\kappa_e = 2.0 \cdot 10^{-16}$ (High),<br>$\kappa_e = 2.0 \cdot 10^{-15}$ (Very high). | 3 |

In the model variation simulations, we implemented the changes to the material parameters on a single brain, the average control brain. Each model variation is listed in table 4, and is explained briefly in the following paragraphs.

#### Variation 1: Pericapillary channel width variations

Due to their small size and their location, the width of capillary PVS is debated. In these simulations we assessed how changes to pericapillary permeability and porosity changed the flow patterns in the brain before, during and after infusion. Using equation 8 from Vinje et al. [2020], we found the pericapillary resistance predicted by the image of Pizzo et al. (2018) to be *R*_*pc*_ = 3.32 *·* 10^−4^ mmHg/ml/min. Two additional possibilities for pericapillary resistances is discussed by Vinje et al. (2020), namely a high resistance scenario based on Bedussi et al. [2018] which suggest a *R*_*pc*_ = 32.24 mmHg/ml/min, and a low resistance scenario where the resistance is computed to be *R*_*pc*_ = 9.2 *·* 10^−3^ mmHg/ml/min. In this model variation all three possibilities were investigated. **Variation 2: Altered Outflow Resistance**. Following the sensitivity analysis on the *R*_out_ parameter performed by Vinje et al. (2020), we ran two simulations where we either doubled (high resistance scenario) or halved (low resistance scenario) the outflow resistance *R*_*DS*_ in equation 10

#### Variation 3: Changes to capillary filtration

Instead of modelling capillary filtration using the compartmental pressure difference between the capillaries and the capillary PVS, we set a constant capillary filtration rate. Vinje et al. (2020) decided on a filtration rate of of 0.16 ml/min in their simulations of constant capillary filtration, a rate we used as well.

#### Variation 4: Altered parenchymal CSF pathways

The exact magnitude and pathway of cerebral CSF flow is uncertain, due to both the small length scales of the cerebral microvasculature Pizzo et al. [2018], and the possible large intrapersonal Xie et al. [2013] and interpersonal Lindstrøm et al. 2019] variations. These uncertainties allow for a certain degree of freedom when it comes to determining flow pathways and transfer coefficients. The simulations in this model variation investigated four scenarios with different relative magnitudes in the transfer coefficients. The changes to each transfer coefficient and their baseline values, with their corresponding case number, is shown in table 4.

#### Variation 5: The importance of extracellular permeability

There exist a great deal of uncertainty concerning the permeability of the extracellular space of the brain. Estimates range from *κ*_*e*_ on the order of 10^−17^ m^2^ Holter et al. [2017] to 10^−15^ m^2^ Ray et al. [2019]. In this model variation, we aimed to investigate how a change in this parameter affect the flow of CSF and ISF, as well as if the response of the human brain to an infusion test changes with extracellular permeability. We have, in addition to the base model of *κ*_*e*_ = 20 nm^2^ also tried a permeability of 10 and 100 times this value.

## Results

### The Brain Stabilises Quickly to Equilibrium

The MPET model with subject specific geometries and generic boundary conditions (see table 4) resulted in interstitial fluid pore speed of 0.7 nm/s before the start of infusion in the control group and 0.8 nm/s in the iNPH group (figure 3, (C)). In both groups, the average fluid pressure in the extracellular space was 10.2 mmHg before the onset of infusion, and the fluid exchange rate between the ECS and both the arterial and venous PVS is less than 0.01 ml/min. The extracellular pressure and fluid speed is shown on the left in figure 3. Right after the start of the infusion, the ISF speed fell in all subjects, before quickly rising and stabilising at an average speed of 1.8 nm/s in the iNPH group and 1.7 nm/s in the control group (figure 3, (C)). The stablisation time was typically about 20 minutes from the onset of infusion, and can be seen on the left hand side in figure 3. Before the onset of infusion, CSF flow in the perivascular spaces was dominated by capillary filtration (data not shown), with a low CSF inflow rate in all subjects from the SAS to the arterial PVS, and a subsequent low through flow in terms of *F*_*pa,pc*_, *F*_*pc,pv*_, *F*_*pa,e*_ and *F*_*e,pv*_. After infusion started, however, fluid transfer in the PVS was dominated by inflow from the SAS and ECS through-flow increased to 0.1 ml/min in both groups, as shown in (figure 3, (E)). Blood flow remained almost constant throughout the entire simulation, shown in (figure 3, (F)).

**Figure 3.**
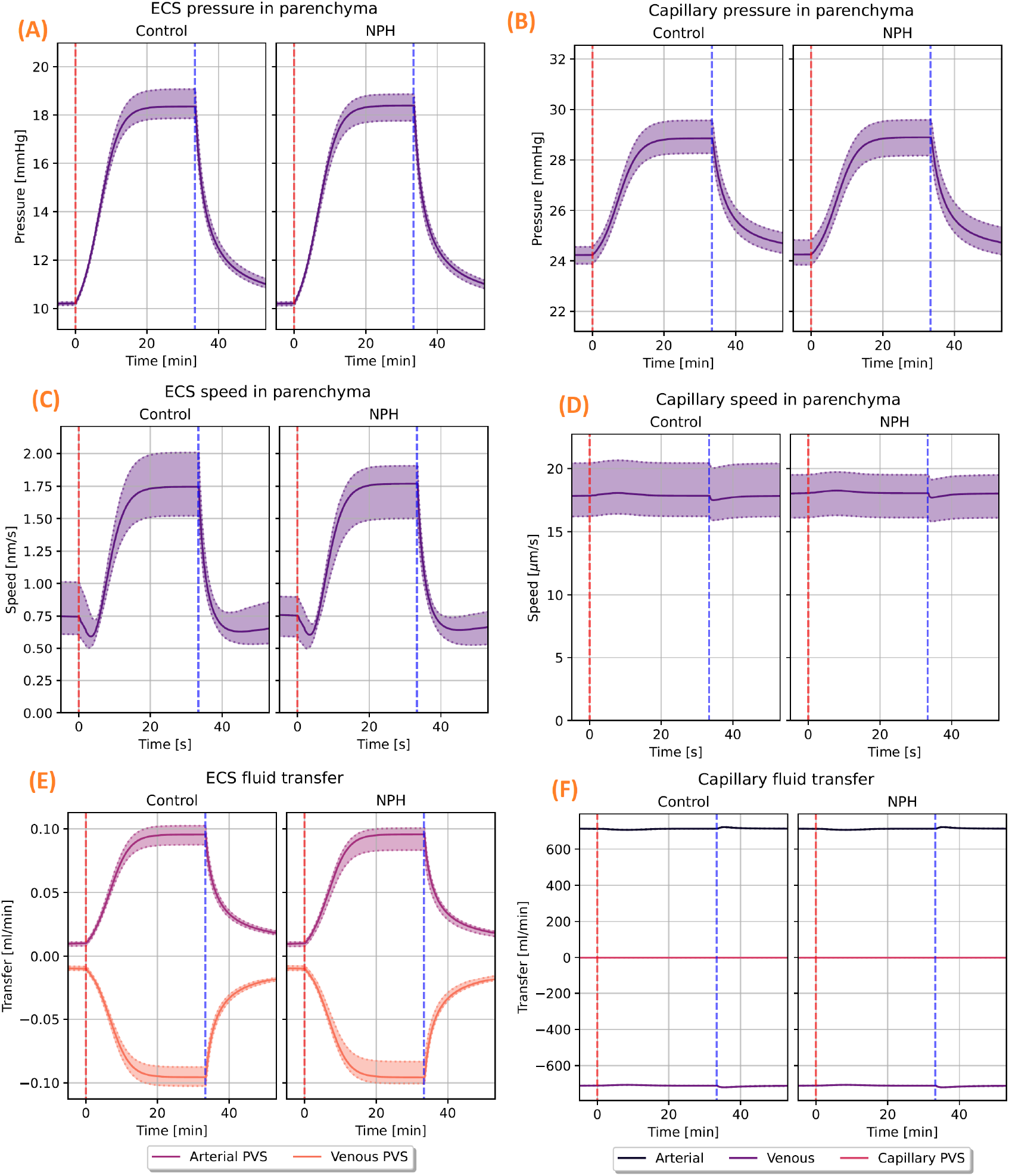
The computed pressure, fluid speed and fluid transfer rates in the ECS and cerebral capillaries. Patient specific geometry and generic boundary conditions were used. On the top row, the computed volume averaged pressure in the ECS (A) and capillaries (B) is shown for both the control and iNPH groups. The middle row shows the corresponding average speed for the ISF in the ECS (C) and capillaries (D). Infusion starts right after *t* = 0 min, marked by the red dashed line,and ends at *t* = 35 min, marked by the vertical blue line. The figures at the bottom row show the net fluid exchange between either the ECS ((E)) or the capillaries ((F)) and their respective connected compartments. A positive value corresponds to a net inflow.

### Base Model 1 - Brain geometry do not explain increased ICP in iNPH during infusion

The difference in how the brain responded to an infusion test between the control and iNPH groups were found to be small when using generic boundary conditions. The total CSF inflow to the ECS from the arterial PVS was 0.1 ml/min in both groups, which is shown in (figure 3, (E)). The largest observed difference in fluid speed between the groups was a 5.9 % increase in the arterial PVS velocity in the iNPH group relative to the control group (data not shown). The relative difference in average pressure between the groups was bounded by 0.9 % which was reached in the venous fluid pressure.

### Base model 2 - Intracranial pressure differences differentiates iNPH patients from controls during infusion

The difference between the average iNPH patient and control increased after subject specific boundary conditions, rather than generic boundary conditions, were applied. The computed volume averaged fluid speed and pressure in the ECS for both groups is shown in figure 4. Within each of the CSF filled compartments, the average peak fluid pressure was higher in the iNPH group than in the control group during infusion. This corresponds to a pressure increase of 21-22% in the iNPH case. The venous blood pressure was 1.5 mmHg (12 %) higher in the iNPH group at the end of infusion compared to the control group, while the arterial blood pressure was only 2 % higher in the iNPH group.

**Figure 4.**
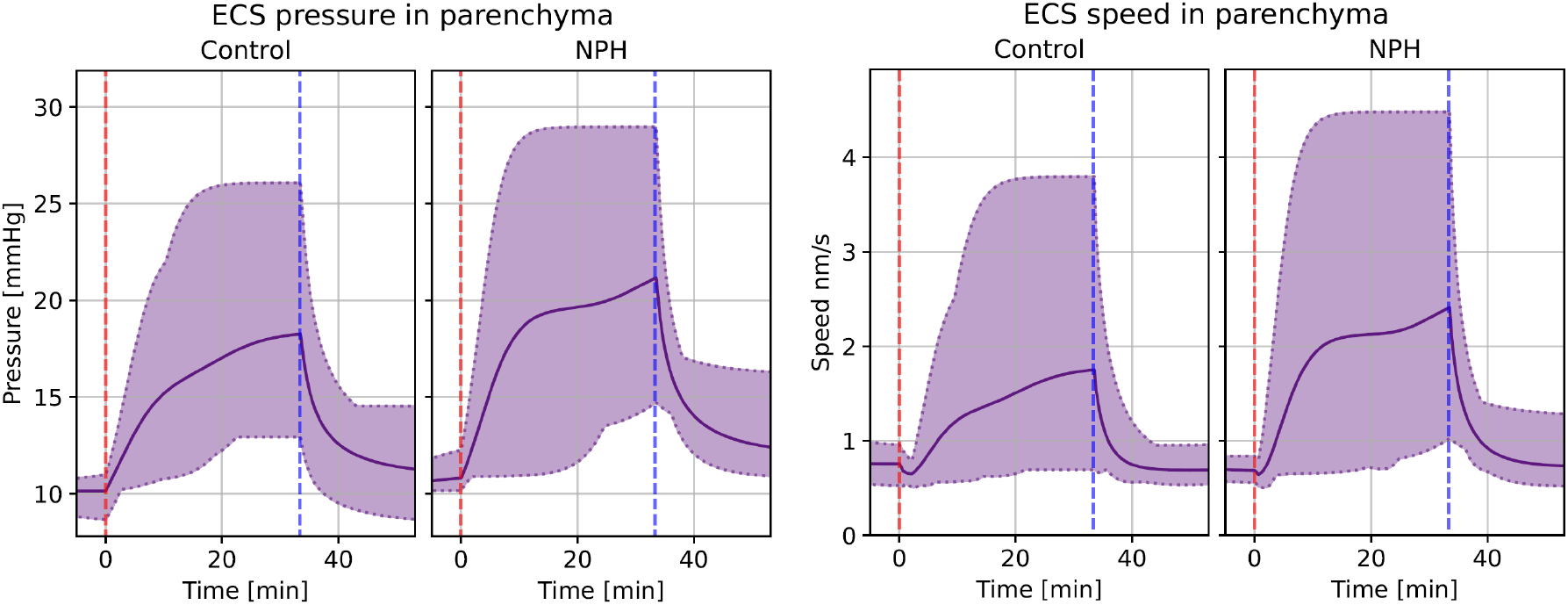
Volume averaged fluid pressure (left) and fluid speed (right) in the extracellular space for both the control and iNPH groups. The group average is shown in a whole line, while the dotted lines show the largest and smallest computed values for the pressure at any given time point. The red dashed line shows the start of infusion, while the blue dashed line marks the end of infusion. These curves show the simulation results when subject specific geometries and boundary conditions were enforced.

The fluid speed within the perivascular compartments and the ECS was on average between 25% (in the capillary PVS) and 75 % larger (in the arterial PVS) in the iNPH group than in the control group (data not shown). The peak ISF speed in the ECS was 2.4 nm/s for the iNPH group compared to 1.8 nm/s for the control group (figure 4). Throughout the entire infusion test, the relative difference between the two groups were always less than 56 %. The relative difference in blood speed was lower than that of CSF speed, reaching at most 6.4 % in the capillaries, with a slightly higher blood speed in the control group (data not shown).

The total CSF flow through the brain was elevated at the end of infusion in the iNPH group. The average CSF flow from the SAS through the brain was 0.70 ml/min in the iNPH group and 0.53 ml/min in the control group, ie 28% and 38% of the combined volume of infusion and production flows through the glymphatic system of the healthy and iNPH patients, respectively. A fifth of this volume went through the ECS, reaching flow rates of 0.09 ml/min in the control group and 0.12 ml/min in the iNPH group at the end of infusion. The capillary filtration rate was nearly equal in the two groups at 0.12 ml/min and 0.11 ml/min in the control and iNPH groups respectively. A full overview over the flow patterns is shown in figure 6. Before the onset of infusion, CSF flow in the perivascular spaces was dominated by capillary filtration, shown in figure 5. In the control group, no CSF enters the arterial PVS at rest, while the total inflow rate for the iNPH group was 0.04 ml/min.

**Figure 5.**
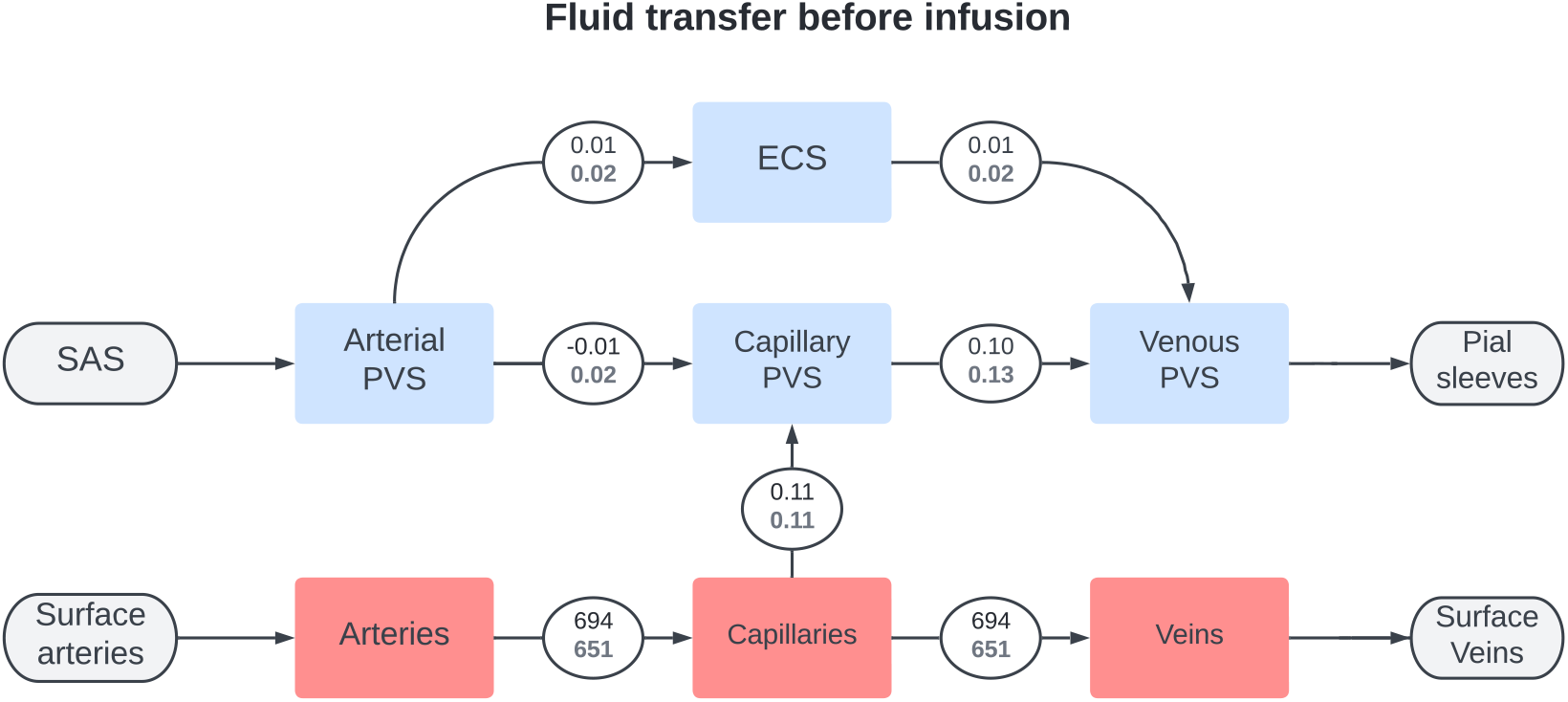
Flowchart showing the total inter-compartmental fluid flow right before the start of infusion (*t* = −1 min). The average fluid transfer rate in ml/min is shown in each connecting circle, with the upper black number being the average in the control group and the lower grey being the iNPH group. These numbers were computed using subject specific geometries and boundary conditions.

**Figure 6.**
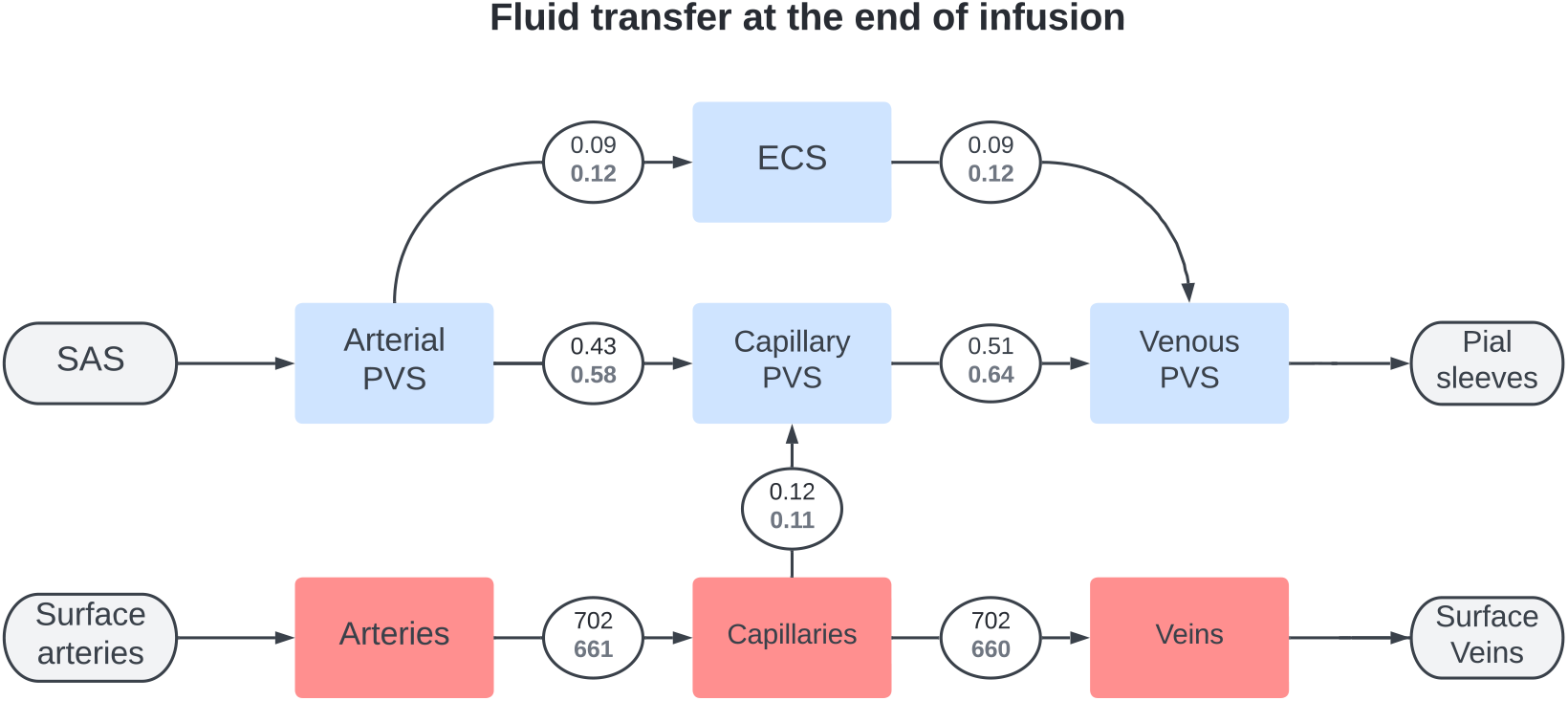
Flowchart showing the total inter-compartmental fluid flow at the end of infusion (*t* = 50 min (3000 s)). The average fluid transfer rate in ml/min is shown in each connecting circle, with the upper black number being the average in the control group and the lower grey being the iNPH group. These numbers were computed using subject specific geometries and boundary conditions.

### Base model 3 - A generic brain geometry gives a reasonable approximation of mean flow in both ECS and PVS

For each subject, we performed one simulation with subject specific boundary conditions and subject specific geometry, and one simulation with subject specific boundary conditions and average geometry. For most subjects, the relative difference in pressure between the two simulations was largest close to the end of infusion for all compartments (figure 7(A)). Fluid speed in every compartment reached a maximal relative difference closer to the start of infusion (figure 7(B)).

**Figure 7.**
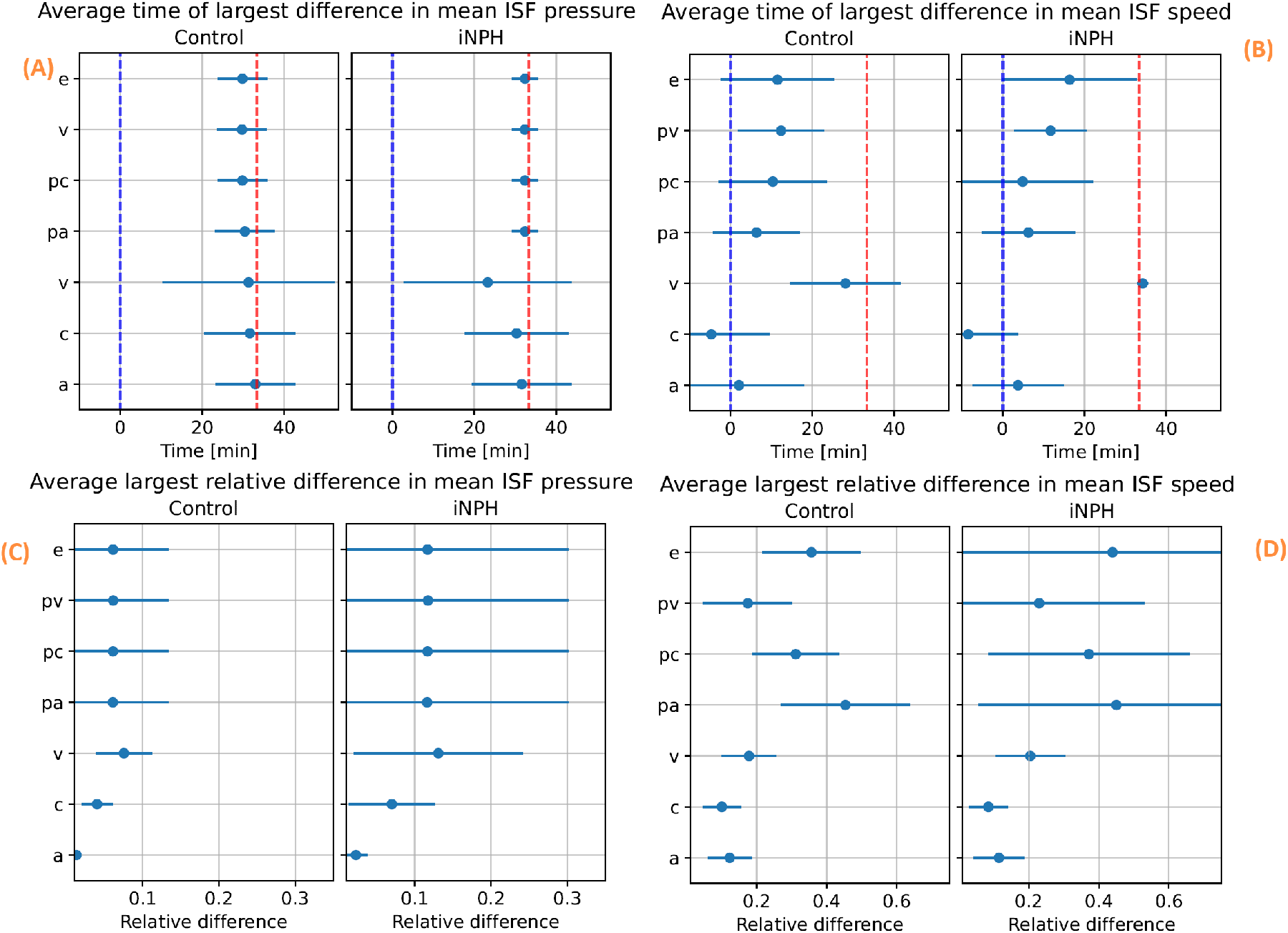
Time averaged relative difference between simulations performed on subject specific geometries and a generic geometry. For each subject, we ran a simulation using the subject specific boundary conditions on both the subject specific and group averaged geometries. Then, for each subject, we computed the relative difference in volume averaged fluid pressure (left) and fluid pore speed (right) between the simulation result when using a subject specific geometry and an average geometry. The top two figures, (A) and (B) show the average time point for the largest relative difference in pressure (A) and pore speed (B) occurs. On the bottom row, the average largest relative difference achieved in terms of both pressure (C) and pore speed (D) in each compartment and group is shown. In all figures, the compartmental group average is denoted with a blue dot, and the lines show the interval defined as the average *±* one standard deviation. In figures (A) and (B) the start of infusion is marked by a blue dashed line, while the end of infusion is marked by a dashed red line.

The difference between subject specific geometries and average geometries in terms of mean pressure was, on average, slightly less than 10 % different in the control group and slightly higher than 10% in the iNPH group. One notable exception, shown at the bottom of (figure 7,(C)), was the arterial blood pressure, where the relative difference was less than 1 % in both groups. Unlike the control group, there was a large intragroup variation among the iNPH patients in the PVS and ECS between the subject specific and average geometries. Here, the standard deviation was almost 0.2, and twice the size of the mean relative difference of 0.1.

In all compartments the fluid speed is more sensitive than the fluid pressure with respect to the brain geometry variations. The largest relative difference in speed between a simulation with a subject specific geometry and average geometry was found in the arterial PVS. Here, the relative difference reached 40 % with a standard deviation of *±* 10% in the control group and *±* 20 % in the iNPH group (figure 7, (D)). There was no discernible trend across the different compartments for when the speed difference is at its largest. In the arterial and capillary compartments the largest difference was reached before the start of infusion, while the time of greatest difference occurred at different times during infusion for the other compartments. The latest point of largest difference happened in the extracellular and venous compartments, close to 20 minutes after the onset of infusion.

### Changes in material parameters yield a changed cerebral waterscape

#### Variation 1 - Uncertainties in pericapillary channel width yields large variations in CSF flow

The choice of pericapillary permeability had little effect on fluid pressure in the ECS, reaching an average of 19.1 mmHg in the high resistance scenario and 18.9 mmHg when using the resistance computed based on the image from Pizzo et al. Pizzo et al. [2018]. With the base model, the ISF pressure reached 18.9 mmHg at the end of infusion. figure 8 shows large variations in mean fluid speed. The pressure values are shown visually at the left side of figure 11. ISF speeds rose with increased flow resistance in the capillary PVS. In the high resistance scenario, fluid in the ECS reached an average speed of 3.7 nm/s, compared to 1.4 nm/s in the base model and 1.1 nm/s in the low resistance scenario (data not shown). The largest difference observed, however, occurs in the capillary PVS where the fluid plateau speed reached 3.9 nm/s in the high resistance scenario and 3300 nm/s in the lowest resistance scenario, shown in figure 8.

**Figure 8.**
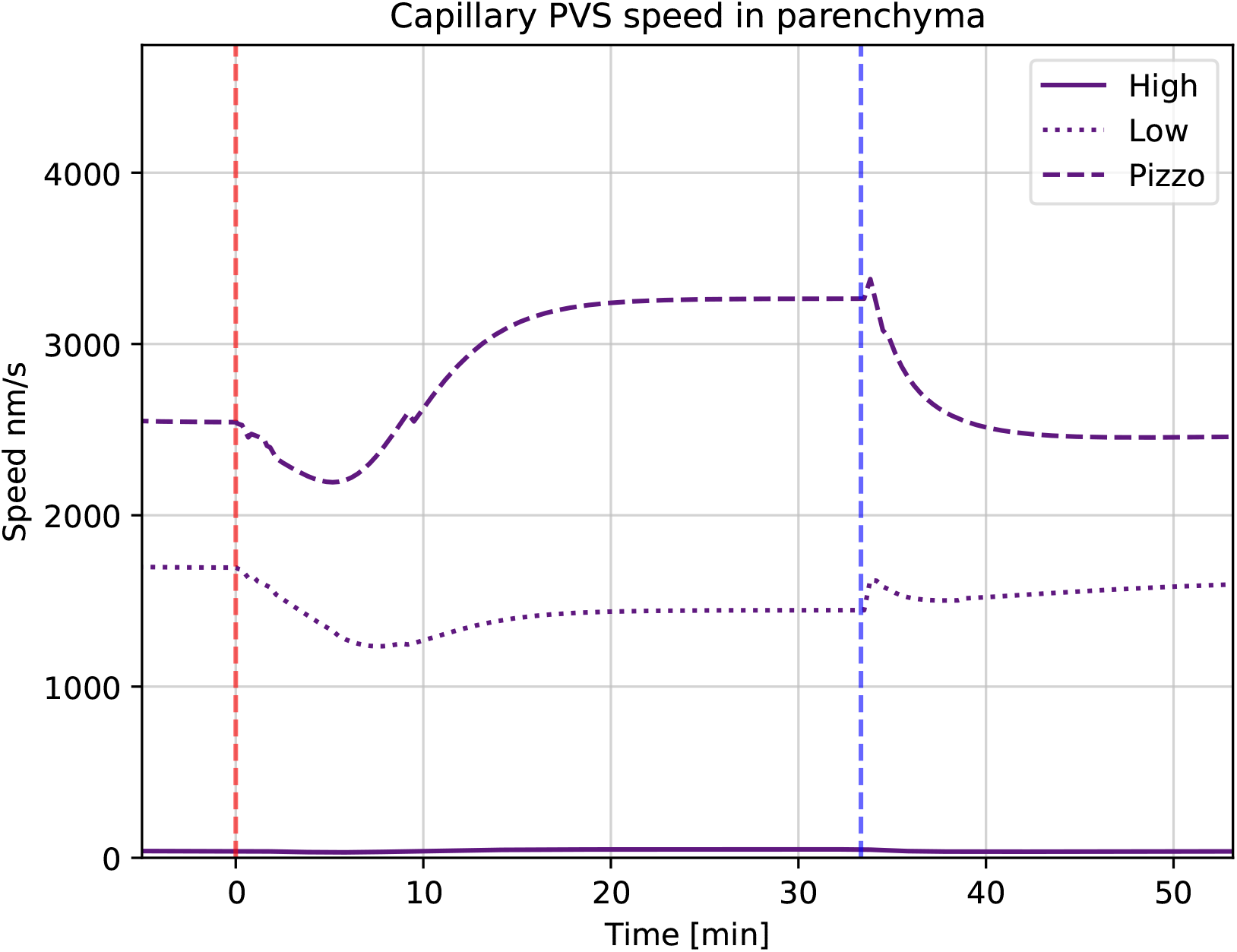
Average fluid speed in the entire capillary PVS as a function of time for each of the three different variations in pericapillary channel width. The red dashed line marks the start of, and the blue dashed line the end of infusion. Here “Low” indicates the base model.

#### Variation 2 - Changes in outflow resistance affect parenchymal CSF flow

A doubled resistance for CSF clearance through the arachnoid granulations resulted in an ISF pressure increase at the end of infusion of 4.2 mmHg on average within the parenchyma, from a plateau of 18.9 mmHg in the base model to 23.1 mmHg in the high resistance scenario. In the low resistance scenario, where the outflow resistance through the arachnoid villi was halved, the pressure plateau fell to 15.2 mmHg, shown in figure 11. Fluid speed in the ECS was similarly affected, with an increase of 0.6 nm/s to 2.0 nm/s in the high resistance scenario and decrease by 0.5 nm/s to a peak average speed of 0.9 nm/s in the low resistance scenario.

#### Variation 3 - Constant capillary filtration yields higher flow rates

A constant capillary filtration rate leads to an increase in fluid pressure and fluid speed in the ECS, shown in figure 11 and figure 12. The ISF pressure at the end of infusion increased from 18.9 mmHg in the base model to 21.0 mmHg, and the average fluid speed is increased from 1.4 nm/s to 4.6 nm/s. While the pressure in the grey matter was lower here than in the base model, an elevated white matter pressure was found close to the lateral ventricles, shown in figure 9. These result show an increased transmantle pressure gradient, defined here as the pressure difference between the ventricular walls and the cortex, going from 0.6 mmHg in the dynamic case to 9.9 mmHg for a constant capillary filtration.

**Figure 9.**
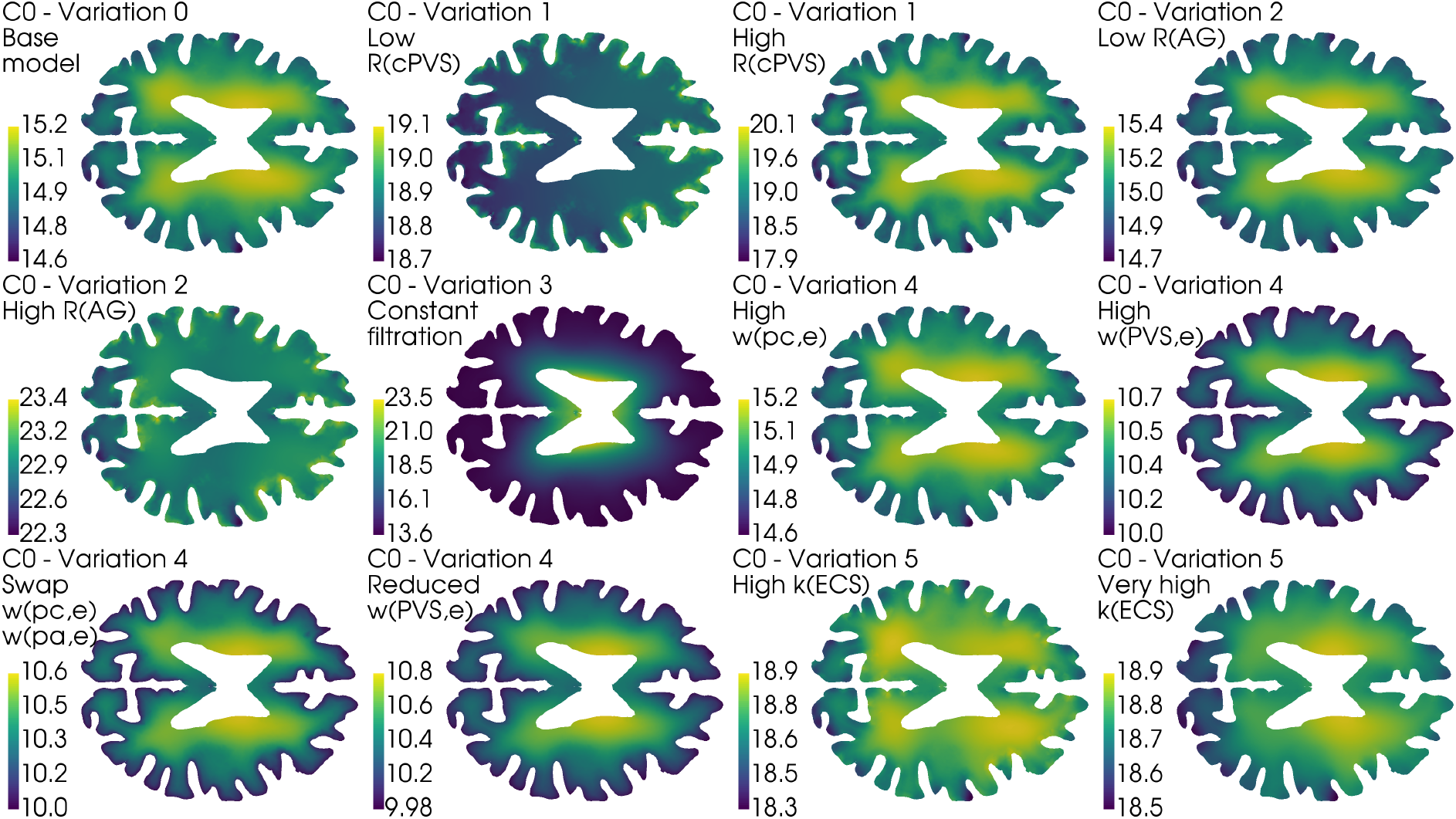
Axial slice of the average control brain geometry with the extracellular pressure field for each of the different model variations in table 4. The base model is shown in the upper left, and serves as a point of comparison with every other result in this figure. The pressure field is computed at the end of infusion, and the pressure values in mmHg is shown in the colourbar to the left of each brain. All cases reveal pressure gradients within the parenchyma, and in the most extreme case (Variation 3, Constant filtration) the transmantle pressure difference is 9.9 mmHg. In most cases (except for Variation 1, Low *R*_cPVS_) the ISF pressure is largest close to the lateral ventricles. This ensures the ISF flow is mainly directed outwards from the ventricles towards the pial surface.

**Figure 10.**
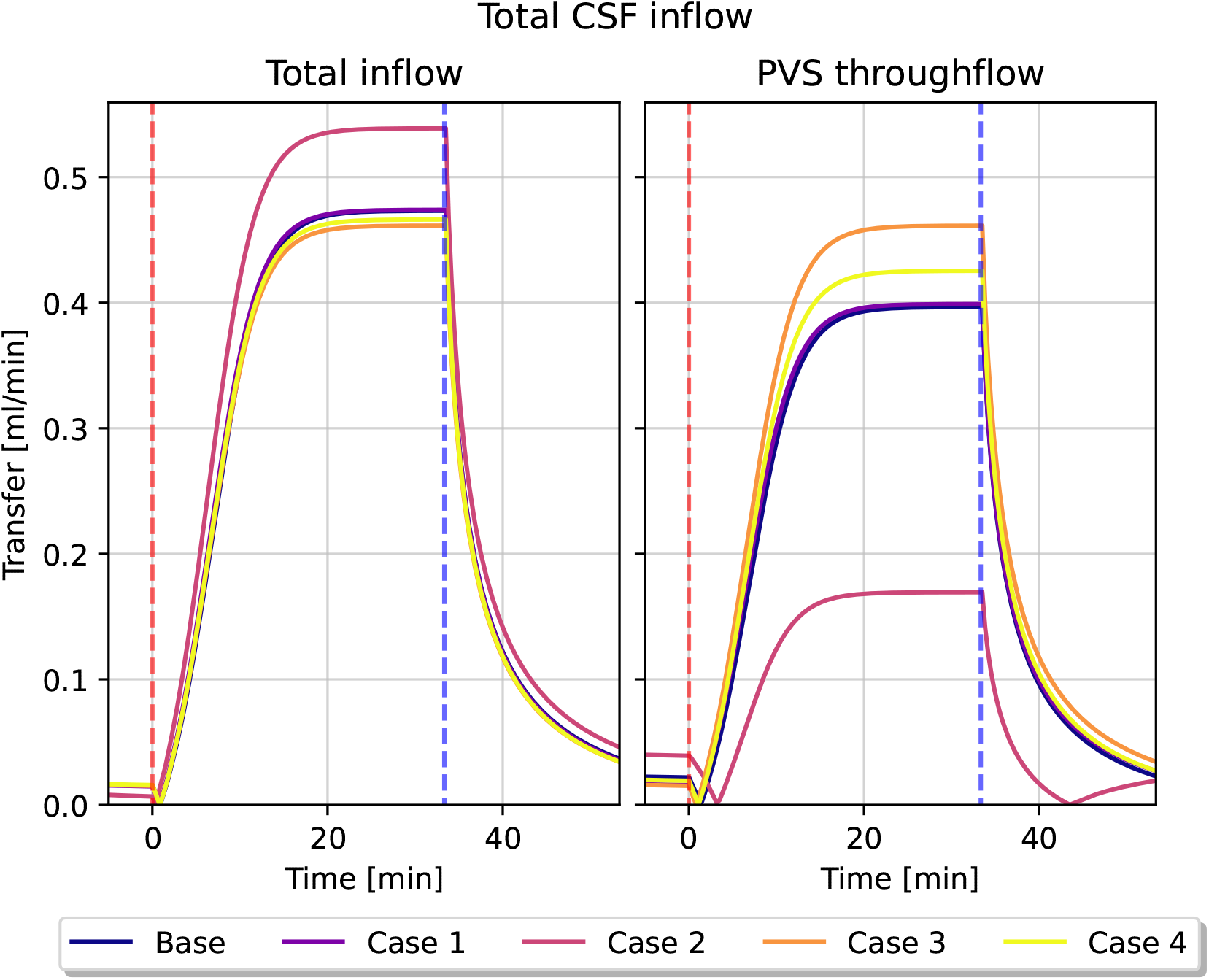
Total CSF inflow, and its chosen pathways for different variations in transfer coefficients. The figure to the left show the total CSF volume that enters the brain through the arterial PVS, while the right figure show how much of this fluid flows into the capillary PVS rather than the ECS. Infusion starts at *t* = 0, and is marked by the red dashed line, and infusion end is marked by the blue dashed line.

**Figure 11.**
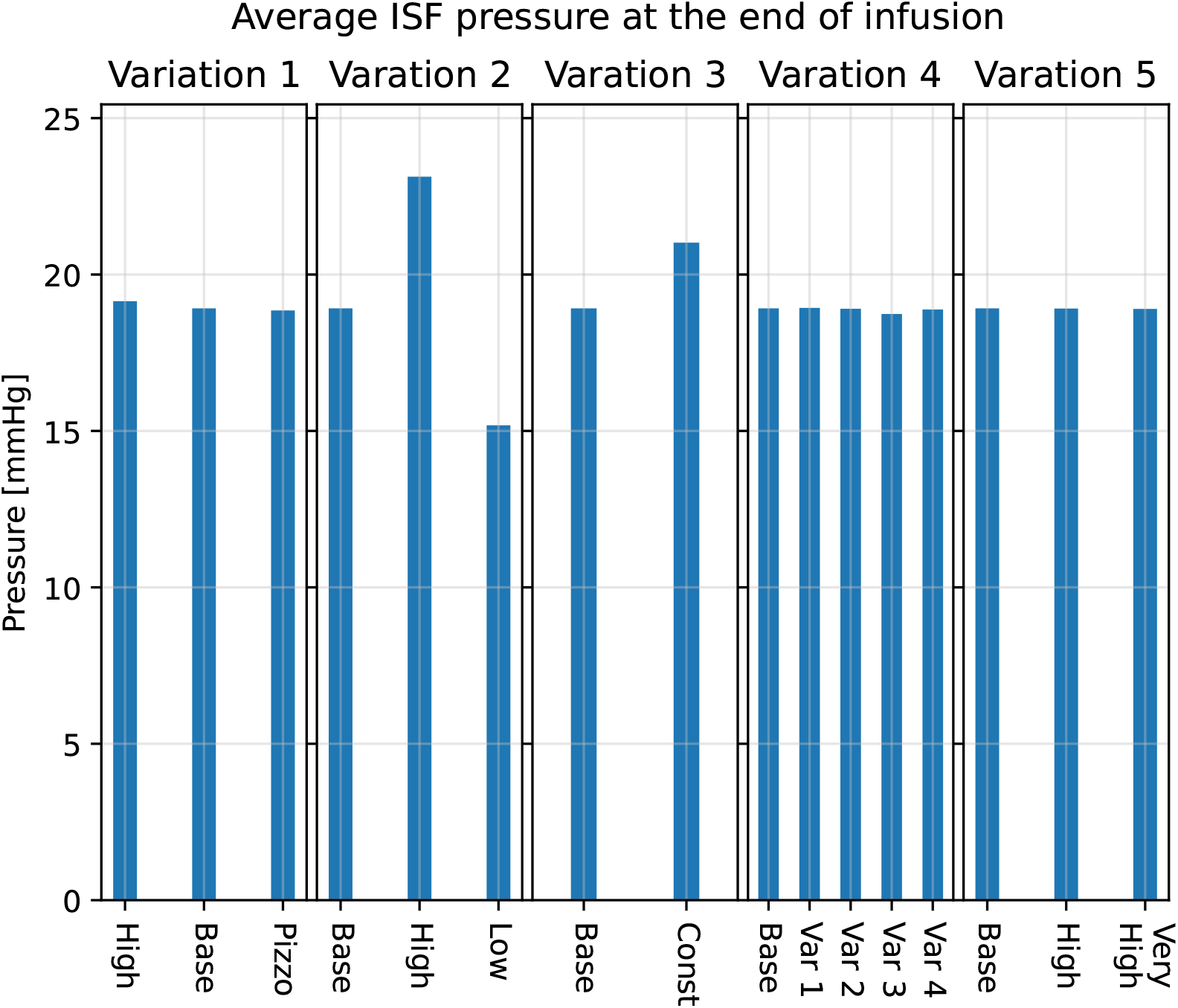
Different model variation effects on average parenchymal fluid pressure in the ECS at the end of infusion (*t* = 50 min (2990 s)). In all figures, base refers to the base model explained in earlier sections. The scale is the same in for all five bar plots. All simulations were performed on the average control group geometry with generic boundary conditions.

**Figure 12.**
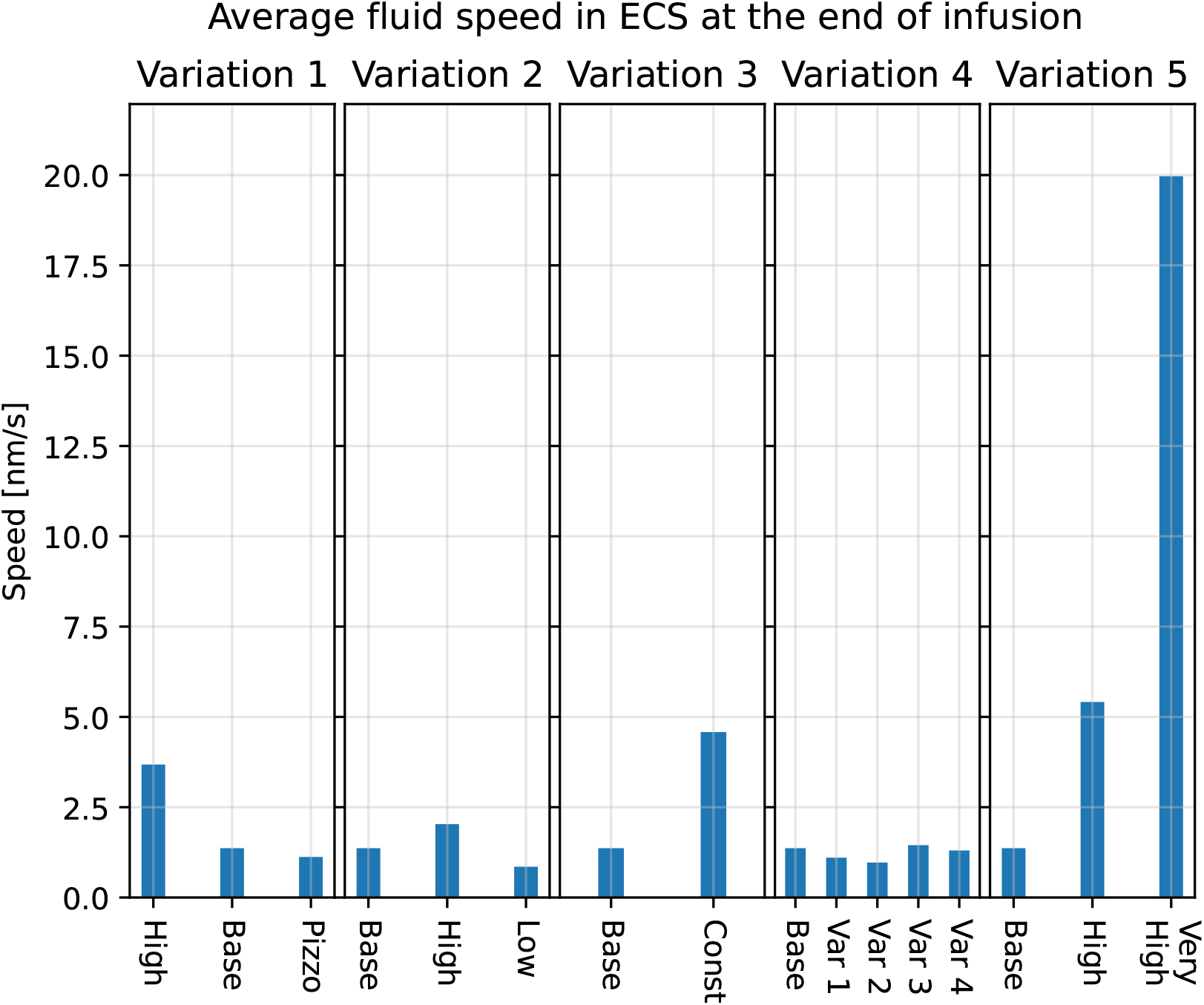
Different model variation effects on average parenchymal fluid speed in the ECS at the end of infusion (*t* = 50 min (2990 s)). In all figures, base refers to the base model explained in earlier sections. The scale is the same in for all five bar plots. All simulations were performed on the average control group geometry with generic boundary conditions.

#### Variation 4 - Changes in transfer coefficients affects total CSF flow through the brain

Fluid transfer patterns between compartments remain similar in most of the transfer coefficient variations. In particular, we found a peak inflow rate of 0.45 ml/min into the arterial PVS in all but one variation. In these cases, fluid flow from the arterial PVS to the capillary PVS were around 0.40 ml/min and 0.45 ml/min, and flow from the arterial PVS to the ECS was limited. In one variation (case 2) the majority of CSF entered the ECS rather than the capillary PVS. The total inflow was 0.55 ml/min, of which less than 0.2 ml/min entered the capillary PVS (figure 10). The average fluid pressure is not affected by the changes in perivascular transfer coefficients, as can be seen in figure 11. The average fluid pressure in the ECS varied by at most 0.2 mmHg from the base model in the transfer coefficient variations. Fluid speed in the ECS remained almost unchanged as the transfer coefficients changed. The base model found a plateau speed of 1.4 nm/s, with the different changes in transfer parameters giving a plateau speed of 1.4 nm/s at the highest and 1.0 nm/s at the lowest. The plateau speeds are shown second from the right in figure 12.

#### Variation 5 - Extracellular permeability changes do not affect parenchymal flow patterns

Changes in extracellular permeability *κ*_*e*_ did not have a large impact on ISF pressure, where all permeability changes tested reached the same plateau pressure of 18.9 mmHg as the base model, shown in the middle graph of figure 11. The ISF speed, however, increased as the permeability rose, going from under 1.4 nm/s in the base model, to 2.0 nm/s in the high permeability scenario and 20.0 nm/s in the very high permeability scenario, shown in figure 12.

## Discussion

### Summary

In our simulation model, when comparing the effect of 1) subject specific brain geometries, 2) subject specific intracranial pressures, and 3) variations in physiological parameters, we can conclude that the variations in physiological parameters cause the largest effects reaching up to a 14-fold (fluid speed). These parameters are also the most uncertain. Next, the variations caused by measured intracranial pressures are up to 75% (fluid speed) and 22% (fluid pressure), while the image based brain geometries cause variations of up to 6% (fluid speed) and 1% (fluid pressure). Finally, we revealed that for differences in ISF pressure to occur, outflow resistance from the SAS (here represented by the combined contribution of *R*_DS_ and *R*_crib_), or capillary filtration close to the ventricular wall may be of particular interest.

### Flow before infusion

The base model, and most model variations, indicate that the flow is not convection-dominated. Before infusion, we found the average ISF pore speed to be less than one nm/s. These speeds are in the same order of magnitude as previously simulated pore speeds Holter et al. [2017], Jin et al. [2016], Dreyer [2022]. The former of these two found that even with a pressure gradient of 1 mmHg/mm, they were not able to get interstitial fluid speeds greater than 1-10 nm/s. In Dreyer [2022], our model was implemented, with some notable differences from the implementation in this article. Here, the authors reported a pre-infusion average speed of 2 nm/s in both groups, corresponding to a pressure gradient of 0.07 mmHg/mm. Our findings with the current model suggest a lower extracellular fluid speed than what was found in Dreyer [2022]. The only major change to the model implemented in Dreyer [2022] and this article is a reduction in pericapillary flow resistance and a inclusion of the *Q*_PVS_ in equation 10. The difference in the magnitude of the ISF fluid velocity suggest a plausible link between cerebrospinal fluid dynamics in the PVS and the rate of flow in the ECS. Our simulations resulted in a pressure average 2.9 mmHg greater in the iNPH group than in the control group at the end of infusion. This pressure difference, indicating the iNPH group has a 22% high peak ISF pressure, is lower than the expected pressure difference in ICP between the two groups. The iNPH group has an outflow resistance *R*_*out*_ which is 80% greater than the control group. This difference in pressure change between the measured CSF pressure and computed ECS pressure might be caused by the inclusion of the *Q*_*pvs*_ sink term in equation 10. This term was included to account for the unphysically large CSF flow rates found in Dreyer [2022]. In Dreyer [2022], the authors, following the results derived by Vinje et al. [2020], excluded this term from the ODE modelling subarachnoid CSF pressure. When the full 3D model of the brain parenchyma was introduced, however, this assumption yielded flow rates of up to 3.0 ml/min through the PVS, more than twice the infusion rate. The *Q*_*pvs*_-term, while successful in bounding the perivascular flux, does also limit the pressure growth on the boundary. It is also worth noting that the pressure difference will, even in the absence of the *Q*_*pvs*_-term still not reach 80%, as the ISF pressure in the iNPH group in Dreyer [2022] is 60 % greater than the control group. Hence, some of this difference between *R*_*out*_ and plateau ISF pressure differences does evidently not stem from the sink term in equation 10, but from other sources.

It can be tempting to suggest enforcing either the infusion pressure directly at the boundary, or to enable a backflow mechanism where (some of) the CSF leaving the venous PVS reenter the SAS, thus increasing the CSF pressure at the boundary. We believe neither of these approaches to be well-advised. The first approach would then inevitably end up in the same situation as Dreyer [2022], as this approach would lack a self-regulating mechanism for bounding the CSF flux through the brain. The second approach, which would necessarily lead to an increased fluid pressure in the SAS, is to our knowledge not quantified and would therefore be speculative, introducing more uncertainty into the model. A third possible approach is to tune the *β*-coefficients in the Robin boundary conditions, but the tests we performed on these parameters did not produce any noticeable effect on the plateau pressure. (Data not shown)

Earlier experimental work has suggested bulk flow velocities in brain tissue of around 0.2 *μ*m/s Rosenberg et al. [1980], Abbott [2004]. In the ECS, ISF pore speeds in the range of 0.58 to 2.50 *μ*m/s has been reported Ray et al. [2019]. These fluid speeds are primarily driven by both a larger pressure gradient than what we found between the arterial and venous PVS, and the assumption of a larger extracellular permeability than the one we used in our model. In a mouse model, Poulain et al. [2023] found average ISF velocities on the order of 3-10 nm/s. Velocities of this magnitude are too small for convection to be the dominating cause of transport of solutes. In our study, we have investigated the effect of changes in extracellular permeability as seen in the middle of figure 12. Even with an increase of extracellular permeability of two orders of magnitude (“Very high” case), the average ISF speed is found to be only 20 nm/s at the end of infusion. Therefore, it is unlikely that the large difference in ISF speed can be explained by the difference in permeability.

In Ray et al. [2019] they manually set a pressure difference between the arterial and venous PVS, and an extracellular permeability. For each set of conditions, they looked at the induced flow in the ECS. To get speeds in the range of 0.58 to 2.50 *μ*m/s, Ray et al. [2019] had to set a pressure of 0.8 mmHg between the arterial and venous PVS. With their chosen distance of 250 *μ*m between the arterial and venous PVS, this pressure difference would lead to a pressure gradient of 3.2 mmHg/mm, significantly larger than pressure gradients in our model. This gradient is in the same magnitude as what Tithof et al. [2022], Boster et al. [2022] found to be necessary to find realistic CSF inflow to the penetrating PVS in mice, at 1.2 - 3.3 mmHg. Yet, as Tithof et al. [2022], Boster et al. [2022] points out, this pressure change is unlikely as the largest possible transmantle pressure gradient in humans is believed to be less than 1 mmHg, see Penn and Linninger [2009]. This limit is further corroborated by Dutta-Roy et al. [2008], whose viscoelastic model predicted the necessary transmantle pressure gradient for the formation of ventriculomegaly to be 1.76 mmHg. Hence, we find it unlikely to be sustained pressure gradients of this magnitude in healthy adults. Our computed average speeds in the ECS correspond to pressure gradients of 0.03 mmHg/mm, and Dreyer [2022] found an average pressure gradient of 0.07 mmHg/mm. Even if the pressure gradients in our study appear small, these gradients are slightly higher than pulsatile transmantle pressure gradients measured experimentally at around 0.0015 mmHg/mm Vinje et al. [2019]. Static pressure gradients over the cerebral aqueduct needed to transport a production of 0.5 L CSF per day has been estimated even lower, at 10^−5^ mmHg/mm Vinje et al. [2019].

### The sensitivity of the brain to infusion tests

The response of the brain to an infusion test have been modelled by several authors, and some articles has also used multi-compartment models. The model by Vinje et al. [2020] served as a baseline for many of our parameter choices, and in many aspects our model agrees with the results of Vinje et al. [2020]. In [Vinje et al., 2020, Fig. 3], the authors show that the intracranial pressure stabilises 20-30 minutes after the onset of infusion, which is in agreement with in vivo infusion tests Kahlon et al. [2005]. Unlike Vinje et al. [2020], we find the amount of CSF entering the brain to increase during an infusion, rather than decrease. This difference might stem from the 1D nature of the model in Vinje et al. [2020], where constant capillary filtration was applied everywhere in the brain. In the present study, we assumed capillary filtration only at the ventricular surface. Regardless of model choice, we find that about a third of the infused CSF enters the brain from the SAS, as shown in figure 10. We remark that it was reported in Vinje et al. [2023] that up to a third of the intrathecal contrast entered the brain.

Poroelastic models have also been used to investigate infusion tests. In Sobey et al. [2012], the authors used a two-compartment poroelastic model to describe the spatial propagation of a sudden increase in ICP in terms of strain and displacement of brain tissue. Notably, they found that even though CSF pressure remained virtually constant throughout the parenchyma, both displacement and strain fell rapidly from the pial surface and into the parenchyma. In most of our model variations, the largest pressure gradients, and hence also fluid velocities are found in the grey matter. This is shown in figure 9, and also documented in [Dreyer, 2022, chp. 7].

The tracer infusion experiments of Iliff et al. [2012], Thrane et al. [2013] and Bedussi et al. [2018] can be regarded as an almost-infusion like scenario due to the relative size between murine and human brains and their infusion rates Vinje et al. [2020]. While direct numerical comparisons might be unfounded, we believe a qualitative comparison might be appropriate. We note, for example that Thrane et al. [2013] found a large extracellular bulk flow at over 1 *μ*m/s in the cerebral extracellular space of mice after injecting tracer directly to the interstitium. Hence, it is clear that bulk flow might be possible if the local pressure gradient is large enough, as might happen during infusion. In alignment with Bedussi et al. [2018] and Iliff et al. [2012], we find the perivascular fluid speed to be significantly higher than interstitial fluid speed in all stages of our simulation. Guo et al. [2020] used MPET to investigate infusion tests and found that the pressure in the PVS and ECS, which in their article was a single combined compartment, should follow the intracranial pressure (pressure in the SAS) during infusion. This finding is supported by our own results, as the pressure graphs in e.g., figure 3 follow the ICP in Vinje et al. [2020], which we used as the model for CSF pressure in the SAS.

The computed pressure fields shown in figure 3 predict that there is little to no difference between healthy individuals and iNPH patients in response to infusion, given that the geometry is the only difference between the groups. In that case, the largest difference between the control and iNPH group, as a group average, is a 5.9 % larger fluid pore speed in the venous PVS. However, in our experimental data, the average *R*_*out*_ in the iNPH group was 18.1 mmHg/(ml/min) compared to 10.0 mmHg/(ml/min) in the control group. With an infusion rate of 1.5 ml/min, this would correspond to a difference of 12.5 mmHg between the two if we assume the response to be linear. The assumption of linearity is unlikely to be true, as the linear relationship between pressure and infusion rate vanish if ICP surpass 26 mmHg Andersson et al. [2008]. Yet, a difference of 6% which is only found in a single quantity of interest within a single compartment, seems too small to be an accurate representation of the groups. Therefore, we find it likely that other parameters than the geometry differ between the groups.

The difference in response between the iNPH and control group grew when subject specific boundary conditions based on the infusion test were implemented. Now, we observe a 21-22% difference in average fluid pressure and between 25% and 75% in fluid speed in all CSF-filled compartments. The difference in ISF pressure and speed is shown in figure 4. The boundary conditions represent the CSF dynamics happening at the surface of our computational domain, ie. the brain surface. However, it is not clear that the lumped parameters of the infusion test can be translated to uniformly distributed boundary conditions in such a simple way as modelled here. In particular, the increased resistance may also express increased resistance of some of the glymphatic pathways within the parenchyma. In particular, our model predicts a positive correlation between *R*_*out*_ and all of our quantities of interest. Figure 4 shows the difference between the control and iNPH groups in terms of ISF pressure and pore speed, and the only difference between the groups is the boundary conditions. Here, both the pressure and fluid speed is higher on average within the iNPH group. In contrast, Eide and Ringstad [2019] discovered that the clearance of CSF tracer from the brain after intrathecal injection was significantly delayed in iNPH patients compared to healthy adults. Furthermore, CSF dynamics and transport in the ventricles differ substantially between the two groups Eide and Ringstad [2019], Eide et al. [2021]. These observations may suggest that our assumption of equal permeability in all compartments in both groups might be incorrect.

While the boundary conditions seem to be important for differentiating between an iNPH patient and a healthy individual, the case for using subject specific geometries is more complicated. The largest relative difference in pressure between an average control and iNPH geometry and a subject specific geometry is found to be around 5% and 10% respectively, as shown in (figure 7, (C)). However, the difference in speed increases to over 20 – 40% in the PVS and ECS in both groups (figure 7, (D)). A possible explanation lies in the fact that the averaging process yielded a smoother and smaller cortical surface. As the largest pressure gradients in our model occur in the grey matter, see either figure 9 or [Dreyer, 2022, chp. 7]. The decreased cortical surface area and relative difference between grey and white matter would yield reduced average pressure and velocity. It is also worth noting that the largest difference in average pressure, shown in (figure 7, (A)), happens at the end of infusion, while the largest difference in fluid speed occurs close to the middle of infusion in all compartments.

Our model included the possibility of flow in capillary PVS. This flow has not been documented experimentally. However, with the approximate size of PVS as shown by Pizzo et al. Pizzo et al. [2018], the capillary PVS resistance is very low, resulting in capillary PVS velocities of around 2-3 *μ*m/s. With all capillary PVS resistances tested in this work, most CSF/ISF flows via capillary PVS rather than the ECS Dreyer [2022]. Particles in these spaces would thus move around 15-20 cm per day, providing a strong clearance mechanism, even without bulk flow in tissue, considering that brain wide clearance occurs on the day scale Ringstad et al. [2018]. If bulk flow occurs only in PVS, diffusion as a transport mechanism is more than sufficient over the small distances from ECS to PVS (25 - 50 *μ*m) over such a time frame Abbott et al. [2010]. However, if velocities in capillary PVS are as high as reported when using the low resistance obtained with data from Pizzo et al. Pizzo et al. [2018], resistances in the arterial and venous PVS might be expected to be lower as well, implying possibly faster clearance rates.

We set a very low transfer coefficient between capillary PVS and the ECS. Hence, our model yields two possible CSF flow pathways, either flowing through through the PVS, or flowing from the arterial PVS to the ECS before reentering the PVS alongside the veins. In most permutations of transfer coefficients in variation 4, the majority of CSF entering the brain through the arterial PVS went to the capillary PVS at the end of infusion, shown in figure 6 and figure 10. The one notable exception, shown in magenta in figure 10 (case 2), is the one where the transfer coefficients between the perivascular compartments and the ECS was increased with a factor 10 (arterial PVS and venous PVS) and a 100 (capillary PVS). This indicates a clear preference for perivascular bulk flow, as suggested by Abbott et al. [2018], rather than the convective flow in the ECS as proposed by, e.g., Iliff et al. [2012], Ray et al. [2019].

When the capillary filtration was applied as a boundary condition at the ventricular wall, rather than a non-zero fluid transfer term between the capillaries and their perivascular spaces, we found significant changes to the cerebral waterscape. The computed transmantle pressure gradient with a constant capillary filtration is found to be 10 mmHg, shown in figure 9. These values are unlikely, as a pressure gradient of this magnitude is significantly larger than the proposed limit of 1.76 mmHg required to form ventriculomegaly Dutta-Roy et al. [2008]. While this pressure gradient is found in the infusion setting, a pressure gradient this large could very well lead to significant damage to brain tissue, something not observed in subjects who have undergone infusion testing.

### Limitations and further work

Our study is to our knowledge the first to model infusion test in a subject specific manner, in the context of the glymphatic system. In order to arrive at such a model we have made a number of simplifications. We considered the CSF dynamics in the subarachnoid space and possible spatial variations in in- and outflow routes to be of minor importance during infusion and that the pressure increases synchronously in the whole subarachnoid space. Furthermore, we have disregarded the spatial deformations and used a scalar valued permeability.

As in Vinje et al. [2020], the majority of the fluid leaves the system along other pathways than the glymphatic pathway (0% at rest and 28% during infusion in the healthy and 12% at rest and 38% during infusion in the iNPH patients, respectively) and the resistance of the pathway is only a fraction of lumped CSF resistance as described by Davson’s equation, cf Andersson et al. [2008] where resistance of 22.3 mmHg/ml/min was established in a group of thirty iNPH patients. We remark that in Vinje et al. [2023] it was observed that about 1/4 of the CSF tracer (gadobutrol) administered by intrathecal injection entered the brain after about 6 hours. To what extent these findings are in conflict or not is not clear and cannot be revealed by the model in this paper as tracer concentrations are not part of the predictions.

Finally, we mention that based on the modeling paper Poulain et al. [2023], we mostly excluded the connection between ECS and capillary PVS in our models. To the authors’ knowledge, the parameter of this pathway is not available in the current literature.

## Conclusion

Infusion tests measure the resistance of the CSF efflux routes and is well established for assessment of iNPH patients. The relationship between this test and the glymphatic system has so far not been adequately modelled or explained. Here, we introduced a subject specific seven compartment model, involving both the vascular, perivascular and extracellular compartments and their interactions under infusion. Surprisingly, we found that the subject specific geometries only play a minor role as compared to the CSF pressure increase under infusion and that other model parameters are more important, such as the infusion pressure, the permeabilities and the transfer coefficients. A considerable amount, but not the majority, of the infused CSF passes through the glymphatic system, according to our computations.

## Declarations

### Author’s contributions

VV, AE, MER, KAM, KHS, LWD developed the mathematical model and designed the study. VV and KHS performed the parameter estimations. VV, LWD performed the simulations. LWD created the figures. AE, SQ, JM performed the infusion tests and MRI imaging. LWD prepared the first manuscript draft. All authors contributed to the editing of the manuscript and figures. All authors read and approved the final manuscript.

## Acknowledgements

We thank Dr. Eric Schmidt, Centre Hospitalier Universitaire de Toulouse France, for discussions and useful insight on infusion testing.

## Ethics approval and consent to participate

Not applicable.

## Consent for publication

Not applicable.

## Availability of data and materials

No original datasets were used in the present study. All parameters used in the model are described in the methods section and illustrated in the supplementary figures. The numerical code and the average control brain geometry is freely available at https://github.com/larswd/MPET-model-iNPH.

Thanks a lot

## References

Jan Malm and Anders Eklund. Idiopathic normal pressure hydrocephalus. Practical Neurology, 6(1):14–27, 2006a.

B Kahlon, G Sundbärg, and S Rehncrona. Lumbar infusion test in normal pressure hydrocephalus. 111(6):379–384, 2005. ISSN 0001-6314.

Ahmed K Toma, Marios C Papadopoulos, Simon Stapleton, Neil D Kitchen, and Laurence D Watkins. Systematic review of the outcome of shunt surgery in idiopathic normal-pressure hydrocephalus. Acta neurochirurgica, 155(10): 1977–1980, 2013.

Sara Qvarlander, Jan Malm, and Anders Eklund. CSF dynamic analysis of a predictive pulsatility-based infusion test for normal pressure hydrocephalus. Medical & biological engineering & computing, 52(1):75–85, 2014.

Anders Eklund, Peter Smielewski, Iain Chambers, Noam Alperin, Jan Malm, Marek Czosnyka, and Anthony Marmarou. Assessment of cerebrospinal fluid outflow resistance. Medical & biological engineering & computing, 45(8):719–735, 2007.

Anders Wåhlin, Khalid Ambarki, Richard Birgander, Noam Alperin, Jan Malm, and Anders Eklund. Assessment of craniospinal pressure-volume indices. American journal of neuroradiology, 31(9):1645–1650, 2010.

Jeffrey J Iliff, Minghuan Wang, Yonghong Liao, Benjamin A Plogg, Weiguo Peng, Georg A Gundersen, Helene Benveniste, G Edward Vates, Rashid Deane, Steven A Goldman, et al. A paravascular pathway facilitates CSF flow through the brain parenchyma and the clearance of interstitial solutes, including amyloid β. Science translational medicine, 4(147):147ra111–147ra111, 2012.

Dennis A Turner. Contrasting metabolic insufficiency in aging and dementia. Aging and disease, 12(4):1081, 2021.

Clifford R Jack Jr, David A Bennett, Kaj Blennow, Maria C Carrillo, Billy Dunn, Samantha Budd Haeberlein, David M Holtzman, William Jagust, Frank Jessen, Jason Karlawish, et al. Nia-aa research framework: toward a biological definition of alzheimer’s disease. Alzheimer’s & Dementia, 14(4):535–562, 2018.

Humberto Mestre, Jeffrey Tithof, Ting Du, Wei Song, Weiguo Peng, Amanda M Sweeney, Genaro Olveda, John H Thomas, Maiken Nedergaard, and Douglas H Kelley. Flow of cerebrospinal fluid is driven by arterial pulsations and is reduced in hypertension. Nature communications, 9(1):4878, 2018.

Aditya Raghunandan, Antonio Ladron-de Guevara, Jeffrey Tithof, Humberto Mestre, Maiken Nedergaard, John H Thomas, and Douglas H Kelley. Bulk flow of cerebrospinal fluid observed in periarterial spaces is not an artifact of injection. bioRxiv, 2020.

N Joan Abbott, Michelle E Pizzo, Jane E Preston, Damir Janigro, and Robert G Thorne. The role of brain barriers in fluid movement in the CNS: is there a ‘glymphatic’ system? Acta neuropathologica, 135(3):387–407, 2018.

Karl Erik Holter, Benjamin Kehlet, Anna Devor, Terrence J Sejnowski, Anders M Dale, Stig W Omholt, Ole Petter Ottersen, Erlend Arnulf Nagelhus, Kent-André Mardal and Klas H Pettersen. Interstitial solute transport in 3D reconstructed neuropil occurs by diffusion rather than bulk flow. Proceedings of the National Academy of Sciences, 114(37):9894–9899, 2017.

Stephen B Hladky and Margery A Barrand. Fluid and ion transfer across the blood–brain and blood–cerebrospinal fluid barriers; a comparative account of mechanisms and roles. Fluids and Barriers of the CNS, 13(1):1–69, 2016.

Stephen B Hladky and Margery A Barrand. The glymphatic hypothesis: the theory and the evidence. Fluids and Barriers of the CNS, 19(1):1–33, 2022.

Lori Ray, Jeffrey J Iliff, and Jeffrey J Heys. Analysis of convective and diffusive transport in the brain interstitium. Fluids and Barriers of the CNS, 16(1):1–18, 2019.

GA Rosenberg, WT Kyner, and E Estrada. Bulk flow of brain interstitial fluid under normal and hyperosmolar conditions. American Journal of Physiology-Renal Physiology, 238(1):F42–F49, 1980.

Erika Kristina Lindstrøm, Geir Ringstad, Angelika Sorteberg, Wilhelm Sorteberg, Kent-Andre Mardal, and Per Kristian Eide. Magnitude and direction of aqueductal cerebrospinal fluid flow: large variations in patients with intracranial aneurysms with or without a previous subarachnoid hemorrhage. Acta neurochirurgica, 161(2):247–256, 2019.

Per Kristian Eide, Lars Magnus Valnes, Erika Kristina Lindstrøm, Kent-Andre Mardal, and Geir Ringstad. Direction and magnitude of cerebrospinal fluid flow vary substantially across central nervous system diseases. Fluids and Barriers of the CNS, 18(1):1–18, 2021.

Per K Eide and Geir Ringstad. Delayed clearance of cerebrospinal fluid tracer from entorhinal cortex in idiopathic normal pressure hydrocephalus: a glymphatic magnetic resonance imaging study. Journal of Cerebral Blood Flow & Metabolism, 39(7):1355–1368, 2019.

Geir Ringstad, Lars M Valnes, Anders M Dale, Are H Pripp, Svein-Are S Vatnehol, Kyrre E Emblem, Kent-Andre Mardal, and Per K Eide. Brain-wide glymphatic enhancement and clearance in humans assessed with mri. JCI insight, 3(13), 2018.

Kerstin Andrén, Carsten Wikkelsø, Per Hellström, Mats Tullberg, and Daniel Jaraj. Early shunt surgery improves survival in idiopathic normal pressure hydrocephalus. European Journal of Neurology, 28(4):1153–1159, 2021. doi:10.1111/ene.14671. URL https://onlinelibrary.wiley.com/doi/abs/10.1111/ene.14671.

Jan Malm and Anders Eklund. Idiopathic normal pressure hydrocephalus. Practical neurology, 6(1):14–27, 2006b. ISSN 1474-7758.

Katie A Peterson, George Savulich, Dan Jackson, Clare Killikelly, John D Pickard, and Barbara J Sahakian. The effect of shunt surgery on neuropsychological performance in normal pressure hydrocephalus: a systematic review and meta-analysis. Journal of neurology, 263(8):1669–1677, 2016. ISSN 0340-5354.

Ian Sobey and Benedikt Wirth. Effect of non-linear permeability in a spherically symmetric model of hydrocephalus. Mathematical Medicine and Biology, 23(4):339–361, 2006.

Benedikt Wirth and Ian Sobey. An axisymmetric and fully 3d poroelastic model for the evolution of hydrocephalus. Mathematical Medicine and Biology: A Journal of the IMA, 23(4):363–388, 2006.

B Tully and Y Ventikos. Cerebral water transport using multiple-network poroelastic theory: application to normal pressure hydrocephalus. Journal of fluid mechanics, 667:188–215, 2011. ISSN 0022-1120.

Tonmoy Dutta-Roy, Adam Wittek, and Karol Miller. Biomechanical modelling of normal pressure hydrocephalus. Journal of biomechanics, 41(10):2263–2271, 2008. ISSN 0021-9290.

Vegard Vinje, Anders Eklund, Kent-Andre Mardal, Marie E Rognes, and Karen-Helene Støverud. Intracranial pressure elevation alters csf clearance pathways. Fluids and Barriers of the CNS, 17:1–19, 2020.

Liwei Guo, John C Vardakis, Dean Chou, and Yiannis Ventikos. A multiple-network poroelastic model for biological systems and application to subject-specific modelling of cerebral fluid transport. International Journal of Engineering Science, 147:103204, 2020.

Jeonghun J Lee, Eleonora Piersanti, K-A Mardal, and Marie E Rognes. A mixed finite element method for nearly incompressible multiple-network poroelasticity. SIAM journal on scientific computing, 41(2):A722–A747, 2019.

Sara Qvarlander, Khalid Ambarki, Anders Wåhlin, Johan Jacobsson, Richard Birgander, Jan Malm, and Anders Eklund. Cerebrospinal fluid and blood flow patterns in idiopathic normal pressure hydrocephalus. Acta neurologica Scandinavica, 135(5):576–584, 2017.

Cees JJ Avezaat and John HM van Eijndhoven. The role of the pulsatile pressure variations in intracranial pressure monitoring. Neurosurgical review, 9:113–120, 1986.

Jan Malm, Johan Jacobsson, Richard Birgander, and Anders Eklund. Reference values for CSF outflow resistance and intracranial pressure in healthy elderly. Neurology, 76(10):903–909, 2011.

Johan Jacobsson, Sara Qvarlander, Anders Eklund, and Jan Malm. Comparison of the csf dynamics between patients with idiopathic normal pressure hydrocephalus and healthy volunteers. Journal of Neurosurgery, 131(4):1018–1023, 2018.

Nina Andersson, Jan Malm, Tomas Bäcklund, and Anders Eklund. Assessment of cerebrospinal fluid outflow conductance using constant-pressure infusion—a method with real time estimation of reliability. Physiological measurement, 26(6):1137, 2005.

John Ashburner. A fast diffeomorphic image registration algorithm. Neuroimage, 38(1):95–113, 2007.

Anders M Dale, Bruce Fischl, and Martin I Sereno. Cortical surface-based analysis: I. segmentation and surface reconstruction. Neuroimage, 9(2):179–194, 1999.

Kent-André Mardal Marie E Rognes, Travis B Thompson, and Lars Magnus Valnes. Mathematical modeling of the human brain: from magnetic resonance images to finite element simulation. Springer Nature, 2022.

Lars-Magnus Valnes and Jakob Schreiner. Svmtk, 2021. URL https://github.com/SVMTK/SVMTK.

Lars Willas Dreyer. Normal pressure with abnormal geometry: A biomechanical model of normal pressure hydrocephalus. Master’s thesis, Department of Mathematics at the University of Oslo, Moltke Moes vei 35, 2022.

Liwei Guo, Zeyan Li, Jinhao Lyu, Yuqian Mei, John C Vardakis, Duanduan Chen, Cong Han, Xin Lou, and Yiannis Ventikos. On the validation of a multiple-network poroelastic model using arterial spin labeling mri data. Frontiers in computational neuroscience, page 60, 2019.

OB Paulson, S Strandgaard, and L Edvinsson. Cerebral autoregulation. Cerebrovascular and brain metabolism reviews, 2(2):161–192, 1990.

Mokhtar Zagzoule and Jean-Pierre Marc-Vergnes. A global mathematical model of the cerebral circulation in man. Journal of biomechanics, 19(12):1015–1022, 1986.

K Kinoshita, A Sakurai, A Utagawa, T Ebihara, M Furukawa, T Moriya, K Okuno, A Yoshitake, E Noda, and K Tanjoh. Importance of cerebral perfusion pressure management using cerebrospinal drainage in severe traumatic brain injury. pages 37–39, 2006.

Paul F Morrison, Douglas W Laske, Hunt Bobo, Edward H Oldfield, and Robert L Dedrick. High-flow microinfusion: tissue penetration and pharmacodynamics. American Journal of Physiology-Regulatory, Integrative and Comparative Physiology, 266(1):R292–R305, 1994.

R Hunt Bobo, Douglas W Laske, Aytac Akbasak, Paul F Morrison, Robert L Dedrick, and Edward H Oldfield. Convection-enhanced delivery of macromolecules in the brain. Proceedings of the National Academy of Sciences, 91 (6):2076–2080, 1994.

Sujit S Prabhu, William C Broaddus, George T Gillies, William G Loudon, Zhi-Jian Chen, and Barlow Smith. Distribution of macromolecular dyes in brain using positive pressure infusion: a model for direct controlled delivery of therapeutic agents. Surgical neurology, 50(4):367–375, 1998.

Peter J Basser. Interstitial pressure, volume, and flow during infusion into brain tissue. Microvascular research, 44(2): 143–165, 1992.

Beatrice Bedussi, Mitra Almasian, Judith de Vos, Ed VanBavel, and Erik NTP Bakker. Paravascular spaces at the brain surface: Low resistance pathways for cerebrospinal fluid flow. Journal of Cerebral Blood Flow & Metabolism, 38(4): 719–726, 2018.

Michelle E Pizzo, Daniel J Wolak, Niyanta N Kumar, Eric Brunette, Christina L Brunnquell, Melanie-Jane Hannocks, N Joan Abbott, M Elizabeth Meyerand, Lydia Sorokin, Danica B Stanimirovic, and Robert G Thorne. Intrathecal antibody distribution in the rat brain: surface diffusion, perivascular transport and osmotic enhancement of delivery. The Journal of physiology, 596(3):445–475, 2018.

Mohammad M Faghih and M Keith Sharp. Is bulk flow plausible in perivascular, paravascular and paravenous channels? Fluids and Barriers of the CNS, 15(1):17, 2018.

Wahbi K El-Bouri and Stephen J Payne. Multi-scale homogenization of blood flow in 3-dimensional human cerebral microvascular networks. Journal of theoretical biology, 380:40–47, 2015.

Hiroshi Ito, Iwao Kanno, Hidehiro Iida, Jun Hatazawa, Eku Shimosegawa, Hajime Tamura, and Toshio Okudera. Arterial fraction of cerebral blood volume in humans measured by positron emission tomography. Annals of nuclear medicine, 15(2):111–116, 2001.

Sang-Pil Lee, Timothy Q Duong, Guang Yang, Costantino Iadecola, and Seong-Gi Kim. Relative changes of cerebral arterial and venous blood volumes during increased cerebral blood flow: implications for bold fmri. Magnetic Resonance in Medicine: An Official Journal of the International Society for Magnetic Resonance in Medicine, 45(5): 791–800, 2001.

Lulu Xie, Hongyi Kang, Qiwu Xu, Michael J Chen, Yonghong Liao, Meenakshisundaram Thiyagarajan, John O’Donnell, Daniel J Christensen, Charles Nicholson, Jeffrey J Iliff, et al. Sleep drives metabolite clearance from the adult brain. science, 342(6156):373–377, 2013.

Adriana T Perles-Barbacaru and Hana Lahrech. A new magnetic resonance imaging method for mapping the cerebral blood volume fraction: The rapid steady-state t 1 method. Journal of Cerebral Blood Flow & Metabolism, 27(3): 618–631, 2007.

Anders Logg, Kent-Andre Mardal, and Garth Wells. Automated solution of differential equations by the finite element method: The FEniCS book, volume 84. Springer Science & Business Media, 2012.

Martin Alnæs, Jan Blechta, Johan Hake, August Johansson, Benjamin Kehlet, Anders Logg, Chris Richardson, Johannes Ring, Marie E Rognes, and Garth N Wells. The fenics project version 1.5. Archive of Numerical Software, 3(100), 2015.

Byung-Ju Jin, Alex J Smith, and Alan S Verkman. Spatial model of convective solute transport in brain extracellular space does not support a “glymphatic” mechanism. The Journal of general physiology, 148(6):489–501, 2016.

N Joan Abbott. Evidence for bulk flow of brain interstitial fluid: significance for physiology and pathology. Neurochemistry international, 45(4):545–552, 2004.

Alexandre Poulain, Jørgen Riseth, and Vegard Vinje. Multi-compartmental model of glymphatic clearance of solutes in brain tissue. Plos one, 18(3):e0280501, 2023.

Jeffrey Tithof, Kimberly AS Boster, Peter AR Bork, Maiken Nedergaard, John H Thomas, and Douglas H Kelley. A network model of glymphatic flow under different experimentally-motivated parametric scenarios. iScience, page 104258, 2022.

Kimberly AS Boster, Jeffrey Tithof, Douglas D Cook, John H Thomas, and Douglas H Kelley. Sensitivity analysis on a network model of glymphatic flow. Journal of the Royal Society Interface, 19(191):20220257, 2022.

Richard D Penn and Andreas Linninger. The physics of hydrocephalus. Pediatric neurosurgery, 45(3):161–174, 2009.

Vegard Vinje, Geir Ringstad, Erika Kristina Lindstrøm, Lars Magnus Valnes, Marie E Rognes, Per Kristian Eide, and Kent-Andre Mardal. Respiratory influence on cerebrospinal fluid flow–a computational study based on long-term intracranial pressure measurements. Scientific reports, 9(1):9732, 2019.

Vegard Vinje, Bastian Zapf, Geir Ringstad, Per Kristian Eide, Marie E Rognes, and Kent-Andre Mardal. Human brain solute transport quantified by glymphatic mri-informed biophysics during sleep and sleep deprivation. Fluids and Barriers of the CNS, 2023.

Ian Sobey, Almut Eisenträger, Benedikt Wirth, and Marek Czosnyka. Simulation of cerebral infusion tests using a poroelastic model. Int. J. Numer. Anal. Model., Ser. B, 3:52–64, 2012.

Vinita Rangroo Thrane, Alexander S Thrane, Benjamin A Plog, Meenakshisundaram Thiyagarajan, Jeffrey J Iliff, Rashid Deane, Erlend A Nagelhus, and Maiken Nedergaard. Paravascular microcirculation facilitates rapid lipid transport and astrocyte signaling in the brain. Scientific reports, 3:2582, 2013.

Nina Andersson, Jan Malm, and Anders Eklund. Dependency of cerebrospinal fluid outflow resistance on intracranial pressure. Journal of neurosurgery, 109(5):918–922, 2008.

N Joan Abbott, Adjanie AK Patabendige, Diana EM Dolman, Siti R Yusof, and David J Begley. Structure and function of the blood–brain barrier. Neurobiology of disease, 37(1):13–25, 2010.

